# Influenza Virus Enhances Bacterial Growth in the Lungs through Weakening Capsule Receptor-Mediated Bacterial Capture by Neutrophils

**DOI:** 10.1101/2024.11.08.622589

**Authors:** Yujie Fang, Wenxia Ma, Chenyun Qian, Jiao Hu, Xianbin Tian, Xueting Huang, Wenlong Lai, Li Wu, Haoran An, Jing-Ren Zhang

## Abstract

Certain serotypes of encapsulated bacteria are more associated with severe pneumonia than others, but the biological reason behind this association has remained largely unknown. This study has uncovered a causal relationship between capsule serotype and virulence phenotype of *Streptococcus pneumoniae*, a leading cause of community-acquired pneumonia, in mice using isogenic bacteria. Certain high-virulence (HV) serotypes persist in the lungs and lead to severe pneumonia, whereas the low-virulence (LV) serotypes are effectively cleared. The capsule type-dependent difference in virulence is determined by the capsule-binding receptors, which recognize the capsular polysaccharides of the LV serotypes (but not the HV serotypes) and enable pathogen capture by neutrophils through complement activation. The capsule receptor-driven immunity of lung neutrophils is significantly compromised by pre-infection of influenza virus, which enhances the survival and virulence of the LV *S. pneumoniae* serotypes, and recurs in the heightened pathogenicity of *Haemophilus influenzae*. This study not only unravels the long-sought mechanistic mystery of serotype-dependent virulence in bacterial lung infections, but also provides an explanation for the well-known pathogenesis synergism between influenza virus and respiratory bacteria.

**Graphical abstract:** 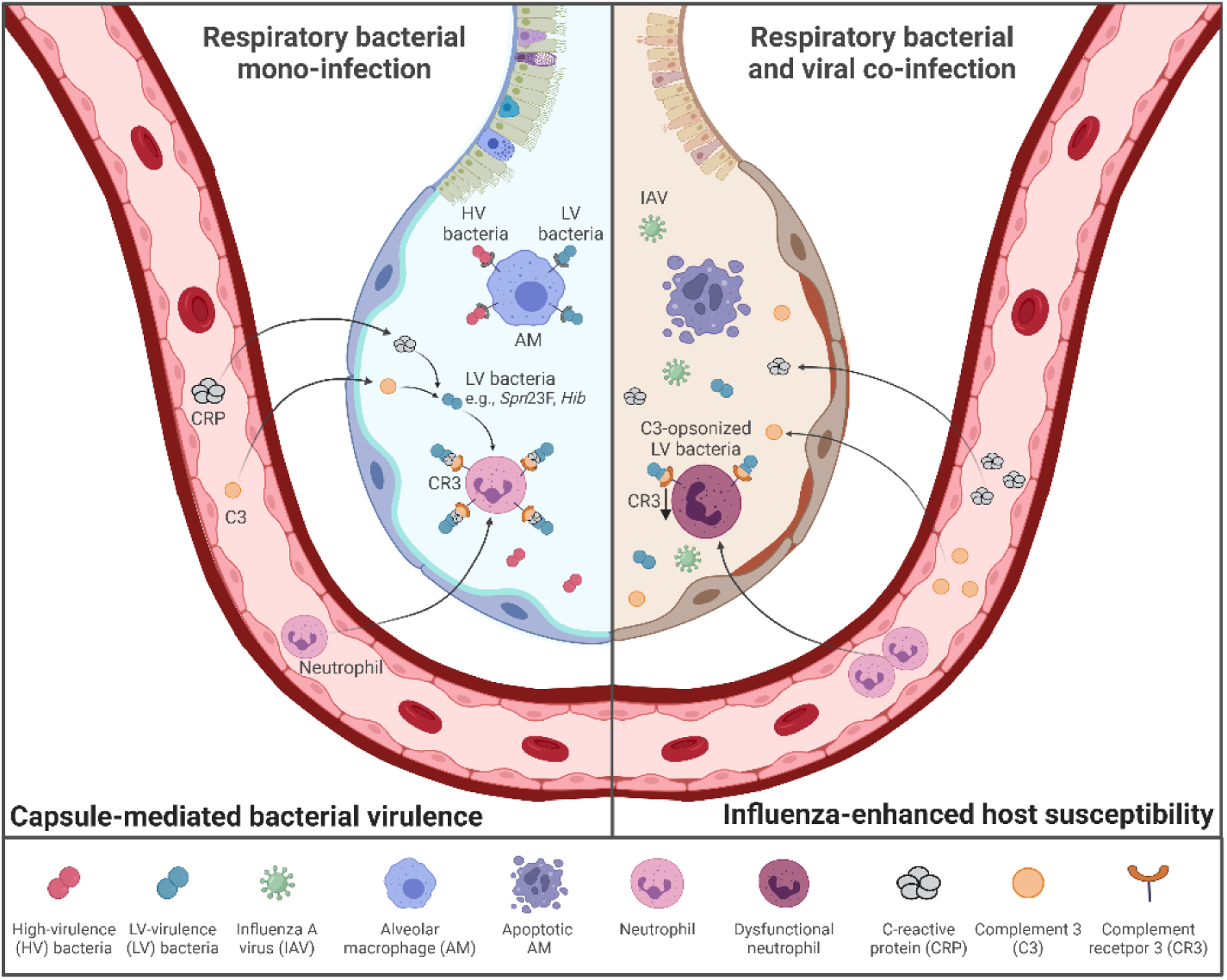

## INTRODUCTION

Encapsulated bacteria, coated by a layer of polysaccharide capsules, represent the most common pathogens of severe pneumonia, the most deadly form of infections.^1^ Capsules share the “slippery” properties that hinder the recognition and binding of immune cells and molecules to encapsulated bacteria.^2^ All polysaccharide capsules are composed of negatively charged repeat units, which, in general, repel the negatively charged mammalian cells.^2,3^ Despite these commonalities, there is a great structural and immunological diversity among capsules. Moreover, many bacteria produce multiple types of structurally and antigenically unique capsules, which are traditionally identified as serotypes by serological reactivity of capsular polysaccharides.^4^ As examples, *Streptococcus pneumoniae* produces at least 104 capsule serotypes;^3,5^ *Haemophilus influenzae* possesses six serotypes of the capsule (serotypes a-f).^6^ Capsule types also determine the pathogenicity potential of encapsulated bacteria, which are manifested by the dominance of the low-numbered pneumococcal serotypes isolated from pneumonia patients,^7–9^ and the overwhelming representation of serotype-b *H. influenzae* in causing invasive infections in young children.^6^

Recent studies have shown that capsules mainly evade molecular recognition and capture by the liver resident macrophage - Kupffer cell (KC) to promote bacterial survival and virulence during septic infections.^10,11^ Moreover, there are great variations among different capsules in the capacity of escaping the hepatic capture, which provide an explanation for the well-known clinical phenomenon - the striking difference in virulence level among capsule types. Based on the differences of capsule types in immune evasion, encapsulated bacteria can be divided into high-virulence (HV) and low-virulence (LV) categories. While the HV serotypes fully bypass hepatic recognition, the LV counterparts are less effective in evading the hepatic immune machinery. More recent studies have uncovered multiple capsule-binding receptors that are responsible for hepatic capture of the LV capsule types, which include the asialoglycoprotein receptor (ASGR) on KCs for serotype-7F and -14 capsules of *S. pneumoniae*,^10^ plasma natural antibodies for serotype-10A and -39 capsules of *S. pneumoniae*,^12^ and plasma C-reactive protein (CRP) for 20 capsule types of *S. pneumoniae*, *H. influenzae* and *Klebsiella pneumoniae*.^13^ However, it remains unknown how capsule types impact the virulence of encapsulated bacteria in the pathogenesis of pneumonia.

*S. pneumoniae* and *H. influenzae* naturally colonize the upper airway of humans as commensals, but they are also the most common pathogens of community-acquired pneumonia and disseminating infections (bacteremia and meningitis).^14,15^ The *S. pneumoniae* and *H. influenzae* diseases share the same seasonal features with those of influenza virus and other respiratory tract viruses, which mostly occur in the winter.^6,16^ The lethal synergism between influenza virus and bacterial co-infections has been observed in the influenza pandemics.^17,18^ In particular, most of the deaths associated with the 1918-1919 influenza pandemic were caused by secondary infections of *S. pneumoniae* and other bacteria.^19,20^ Previous studies have shown that influenza virus-induced interferon-γ suppresses anti-bacterial phagocytosis of alveolar macrophages (AMs).^21^ Pre-infection of influenza virus has also been documented to impair the ion channel activity of lung epithelial cells,^22^ and make lung epithelial cells more prone to bacterial toxin-induce cell death.^23^ It has been well established that neutrophils are rapidly recruited into the lungs during acute pneumonia.^24,25^ However, the precise contributions of neutrophils to anti-bacterial immunity in the lungs remain largely unclear.

In this work, we determined the functional influence of capsule serotypes on immune evasion and virulence of encapsulated bacteria in mouse pneumonia model. Using isogenic strains producing different types of *S. pneumoniae* capsule, we found that capsule types define the high/low-virulence phenotypes in the lung infection. Infiltrating capsule-binding receptors and neutrophils were identified as the major immune effectors that shape the serotype-dependent survival and virulence of *S. pneumoniae* in the lungs, linked through the complement pathway. Influenza-impaired capsule receptor-mediated immunity of lung neutrophils was an important reason for the host hyper-susceptibility to LV *S. pneumoniae* serotypes, further validated by the enhanced pathogenicity of *H. influenzae*.

## RESULTS

### Capsule type defines the virulence level of *S. pneumoniae* in lung infection

Our previous studies have found that capsular serotypes of *S. pneumoniae* define bacterial ability to escape the capture by liver resident macrophage - Kupffer cell and thereby virulence level in septic infection.^10^ While this finding provides an explanation for the serotype-dependent virulence in invasive pneumococcal disease (IPD), the hepatic mechanism cannot explain why certain serotypes are more prevalent than the others in the pneumonia.^26^ We sought to address this question by assessing the virulence level of wild-type *S. pneumoniae* strains representing 13 serotypes in a mouse pneumonia model (**Fig. 1A**). These strains displayed striking variations in virulence post intranasal (i.n.) inoculation of 1 × 10^7^ colony forming units (CFU). While all of the mice infected with serotypes 2, 3, 4, 6A, and 8 died within 7 days, infection with serotypes 6B, 7F, 9V, 14, 18C, 19A, 19F, and 23F did not yield any mortality (**Fig. S1A**). These serotype-specific phenotypes were further verified with capsule-switched strains that expressed one of the 13 different capsules in the same genetic background (TH870, serotype 6A) (**Fig. 1B**). These strains showed similar serotype-dependent high-virulence (HV) and low-virulence (LV) phenotypes as observed in septic infection model,^10^ indicating differences in virulence. The HV serotypes exhibited higher bacterial loads in the lungs at 24 hours post infection (hpi) with an average number of 5 × 10^6^ CFU per lung, which was 132- fold higher than the LV counterparts (**Fig. 1C and S1B**). Moreover, i.n. inoculated HV pneumococci tended to invade into the blood, with 82% of the mice exhibiting bacteremia at 24 hpi; in contrast, most of the mice inoculated with LV strains did not show systemic dissemination (**Fig. 1D and S1C**). These results indicated that capsule types shape pneumococcal virulence in respiratory infection.

**Figure 1.**
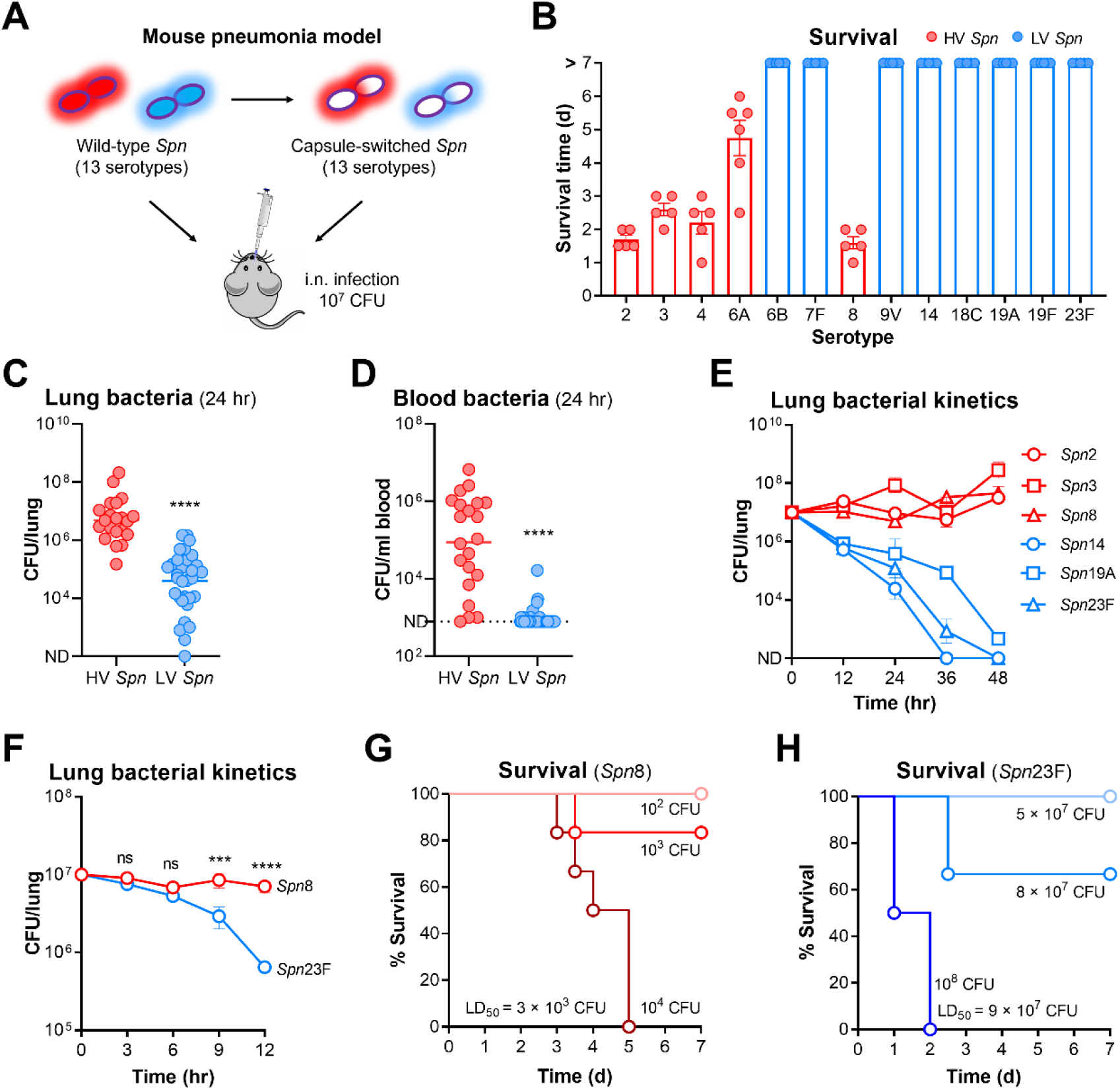
Impact of capsule types on the virulence levels of *S. pneumoniae* in lung infection. **(A)** Schematic workflow of assessment for the virulence level of wild-type and capsule-switched *S. pneumoniae* strains in mouse pneumonia model. **(B)** Survival time of CD1 mice intranasally (i.n.) infected with 10^7^ CFU of capsule-switched *S. pneumoniae* strains. HV and LV serotypes were indicated with red and blue dots, respectively. n = 5-6. **(C and D)** Bacterial loads at 24 hr in the lungs (C) and blood (D) of mice infected as in (B). Data were combined with 3-6 mice per group for each strain, and each dot represented one mouse. **(E)** Bacterial kinetics within 48 hr in the lungs of mice i.n. infected with 10^7^ CFU of HV (serotypes 2, 3, and 8) and LV (serotypes 14, 19A, and 23F) *S. pneumoniae* strains. n = 3-6. **(F)** Bacterial kinetics within 12 hr in the lungs of mice i.n. infected with 10^7^ CFU of HV *Spn*8 and LV *Spn*23F. n = 5-6. **(G and H)** Survival time of mice i.n. infected with 10^2^-10^4^ CFU of HV *Spn*8 (G) and 5-10 × 10^7^ CFU of LV *Spn*23F (H). LD_50_ for each strain was calculated and shown. n = 6. Unpaired *t* test (C, D), ordinary two-way ANOVA with Sidak’s multiple comparisons test (F), ns, not significant, ***, P < 0.001, ****, P < 0.0001.

To define how serotypes modulate pneumococcal virulence in the lung, we examined bacterial dynamics with representative isogenic HV and LV strains. CFU enumeration revealed persistence of HV bacteria (serotypes 2, 3, and 8), but effective eradication of LV variants (serotypes 14, 19A, and 23F) in the lung within 48 hpi (**Fig. 1E**). Mice infected with serotypes 2, 3, and 8 pneumococci developed bacteremia at 24 hpi, which was worsen at 48 hpi. By comparison, no bacteria of serotypes 14, 19A, and 23F were detected in the blood circulation during the same period (**Fig. S1D**). Because the HV and LV serotypes showed clear differences in lung bacteria as early as 12 hr before the onset of bacteremia, it appeared that the virulence phenotypes were shaped in the first 12 hr. We thus characterized the fates of representative HV (serotype 8, *Spn*8) and LV (serotype 23F, *Spn*23F) strains in the early phase of lung infection. While comparable bacterial levels were observed for the two serotypes in the first 6 hr, *Spn*23F started to show significant reduction in the lung at 9 hr and continued to decline afterwards. By comparison, *Spn*8 was consistently maintained at a high level during this period (**Fig. 1F**). This serotype-specific virulence was also manifested by a 30,000-fold difference in 50% lethal dose (LD_50_) between *Spn*8 (3 × 10^3^ CFU) and *Spn*23F (9 × 10^7^ CFU) (**Fig. 1G and 1H**). These observations have demonstrated that the innate immunity in the lungs of mice is much more capable of eliminating the LV pneumococci at the early phase of lung infection.

### Neutrophil immunity defines the serotype-specific virulence phenotypes in the lung

Although a number of immune cells and molecules have been described to contribute to pulmonary defense against pneumococcal infection,^27–29^ it is completely unknown how lung innate immunity discriminates HV and LV *S. pneumoniae* serotypes. We first detected immune cell dynamics in the alveolar space of mice by flow cytometry within 12hr following i.n. inoculation with *Spn*8 and *Spn*23F pneumococci. Alveolar macrophages (AMs) accounted for 88% cellular population in BALF samples from uninfected mice (**Fig. S2A**). Consistent with the apoptotic death of AMs in response to pneumococcal infection,^30^ the cell number of AMs gradually reduced in the first 12 hr of lung infection (**Fig. 2A**). In contrary, neutrophils became the major immune cells at 9 and 12 hr (**Fig. 2A**). Although migrating monocytes have been shown to contribute to pulmonary resistance against pneumococcal infection,^29^ they remained as a minor fraction of immune cells in BALF during the first 12 hr (**Fig. S2A**). This infection stage-dependent shift of AMs and neutrophils in the lungs of mice has been shown in the previous studies that were conducted with HV serotypes 2, 3, and 4.^30–32^

**Figure 2.**
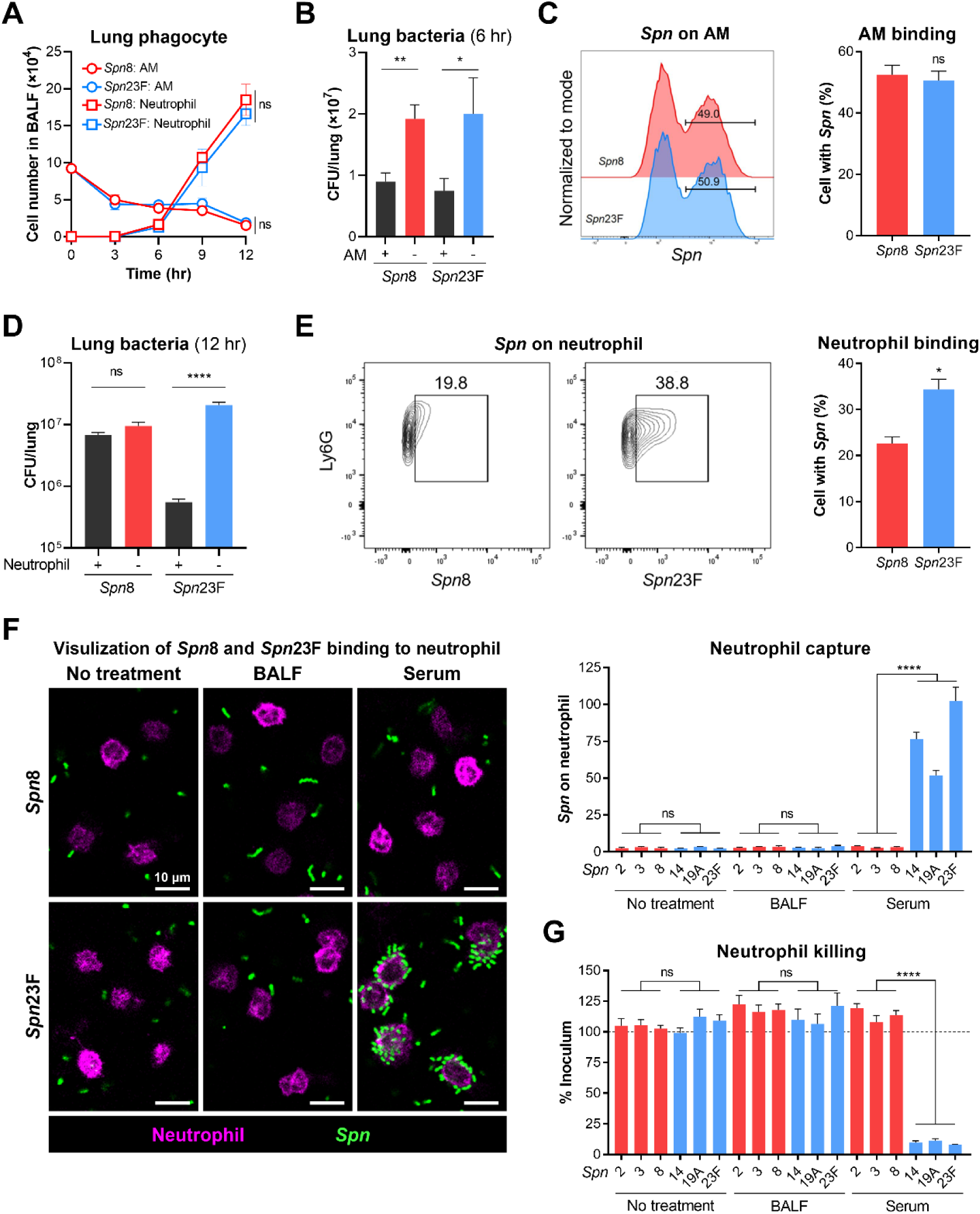
Preferential elimination of *S. pneumoniae* with LV serotypes by neutrophils in lung infection. **(A)** Cell number of alveolar macrophages (AMs) and neutrophils in the BALF of C57BL/6 mice i.n. infected with 10^7^ CFU of *Spn*8 and *Spn*23F. n = 3. **(B)** Bacterial loads at 6 hr in the lungs of WT or AM-depleted mice i.n. infected with 10^7^ CFU of *Spn*8 and *Spn*23F. AMs were depleted by i.n. inhalation with 50 μl of clodronate liposomes (CLL) 5 days before infection. n = 4-6. **(C)** Flow cytometry analysis of pneumococcus-binding AMs at 6 hr in the BALF of WT mice infected as in (A). *S. pneumoniae* strains were labeled with the fluorescence of AF647. n = 3. **(D)** Bacterial loads at 12 hr in the lungs of WT or neutrophil-depleted mice i.n. infected with 10^7^ CFU of *Spn*8 and *Spn*23F. Neutrophils were depleted by i.p. injection of 500 μg anti-Ly6G antibody (clone 1A8) one day before infection and 100 μg 1A8 antibody at the time of infection. n = 6. **(E)** Flow cytometry analysis of pneumococcus-binding neutrophils at 9 hr in the BALF of WT mice infected as in (A). *S. pneumoniae* strains were labeled by expressing the fluorescence protein RFP. n = 3. **(F)** Fluorescent imaging (left) and quantification (right) of *S. pneumoniae* binding to primary neutrophils under the conditions of no treatment, BALF, and normal serum. n = 6 random fields of view (FOVs). Scale bar, 10 μm. **(G)** Killing levels of *S. pneumoniae* by primary neutrophils under the conditions of no treatment, BALF, and normal serum. Killing ratio was calculated by dividing remaining viable bacteria by the inoculum. n = 4-6. Ordinary two-way ANOVA with Sidak’s multiple comparisons test (A), unpaired *t* test (B, C, D, E), ordinary one-way ANOVA with Dunnett’s multiple comparisons test (F, G), ns, not significant, *, P < 0.05, **, P < 0.01, ****, P < 0.0001.

Given that AMs were the predominant immune cells in the lung at the initial stage of pneumococcal infection, we characterized their roles in pneumococcal clearance during this period. We firstly depleted the AMs in mice by inhalation of clodronate liposomes (CLL) prior to i.n. infection. The CLL-mediated AM depletion resulted in a significant increase in lung bacteria at 6 hpi, but there was a similar level of bacterial burden in the lungs of mice infected with *Spn*8 and *Spn*23F (**Fig. 2B**). This finding was further validated with the AM conditionally knockout (AM-cKO) mice.^33^ Similar to what was observed with the CLL-treated mice, the bacterial burdens in the lungs of WT and AM-cKO mice were comparable between the two HV and LV pneumococcal strains (**Fig. S2B**). Moreover, flow cytometry revealed that AMs in the BALF captured a similar degree of *Spn*8 and *Spn*23F (**Fig. 2C**). Finally, we assessed the importance of AMs in host survival during pneumococcal lung infection. In contrast to full survival of WT mice, all the AM-deficient mice showed higher levels of susceptibility to i.n infection of *Spn*8 and *Spn*23F (**Fig. S2C**). These findings revealed that AMs are vital for suppressing pneumococcal growth in the early phase of lung infection, but they do not contribute to the observed differences in virulence between HV and LV serotypes.

Given the abundance of neutrophils in the alveolar space after 9 hr following pneumococcal infection, we evaluated their potential roles in clearing HV and LV serotypes in the lungs by selective depletion with 1A8 antibody. Neutrophil depletion led to dramatic outgrowth of LV *Spn*23F at 9 and 12 hr (**Fig. S2D**), which represented inflammatory stage of lung infection with massive infiltration of neutrophils (**Fig. 2A**). Specifically, the *Spn*23F bacteria in the lungs of 1A8-treated mice were 38-fold higher than those in normal mice at 12 hr. However, the absence of neutrophils did not result in apparent difference in HV *Spn*8 loads in the lung during the first 12 hr (**Fig. 2D**). We further compared the binding ability of neutrophils to HV and LV serotypes under the *in vivo* conditions by identifying neutrophils from BALF of i.n. infected mice at 9 hr. Flow cytometry revealed a significantly higher level of neutrophil binding to LV *Spn*23F, as compared to HV *Spn*8 (**Fig. 2E**). Moreover, neutrophil-depleted mice were highly susceptible to i.n. challenge with 5 × 10^7^ CFU of *Spn*23F, an otherwise non-lethal dose in WT mice, whereas there was no difference in the fatality of mice with or without neutrophils post infection with 10^3^ CFU of *Spn*8 (**Fig. S2E**). Together, our data demonstrated that, at the inflammatory stage of lung infection, infiltrated neutrophils are capable of effective elimination of the LV pneumococci, but fail to control the HV bacteria.

We finally investigated how neutrophils capture the LV pneumococci by using primary neutrophils isolated from the lungs of mice after 12 hr of *Spn*23F infection. Neutrophils showed extremely low binding to both HV and LV *S. pneumoniae* even with supplementation of the BALF supernatant (**Fig. 2F**). However, addition of normal mouse serum dramatically enhanced neutrophil binding to all three representative LV serotypes (14, 19A, and 23F); but the treatment had no impact on neutrophil binding to HV serotypes (2, 3, and 8) (**Fig. 2F**). Consequently, over 90% of the serum-opsonized LV pneumococci were killed by neutrophils after incubation for 30 min, whereas the HV bacteria were completely resistant to the bactericidal activity (**Fig. 2G**). These observations indicated that lung neutrophils depend on a soluble factor(s) in the plasma to capture and kill LV pneumococcal serotypes.

### C-reactive protein enables neutrophils to capture serotype-23F *S. pneumoniae*

To determine if neutrophils directly recognize the capsules of the LV serotypes, we used free capsular polysaccharides of *Spn*23F (CPS23F) to competitively inhibit neutrophil-mediated anti-bacterial activity. As compared with the untreated control, the instillation of CPS23F into the airway with bacterial inoculation resulted in a 7-fold increase in lung bacteria at 12 hr. This CPS blocking effect was serotype-specific because similar treatment with the CPS of serotype-14 *S. pneumoniae* (CPS14), an LV serotype, did not show an obvious effect (**Fig. 3A**). Under the *in vitro* conditions, free CPS23F completely blocked neutrophil binding to and killing of *Spn*23F bacteria (**Fig. 3B and 3C**). The CPS23F-based inhibition against neutrophils was serotype-specific since CPSs of other tested serotypes (8, 14, and 19A) did not affect the immunity of neutrophils to *Spn*23F (**Fig. 3C**). This serotype-specific inhibition of neutrophil clearance by CPS was further supported by parallel assessments with LV serotypes 14 and 19A (**Fig. S3A and S3B**). These results strongly suggested that the capsules are the primary targets for neutrophil capture of LV serotypes in the lung.

**Figure 3.**
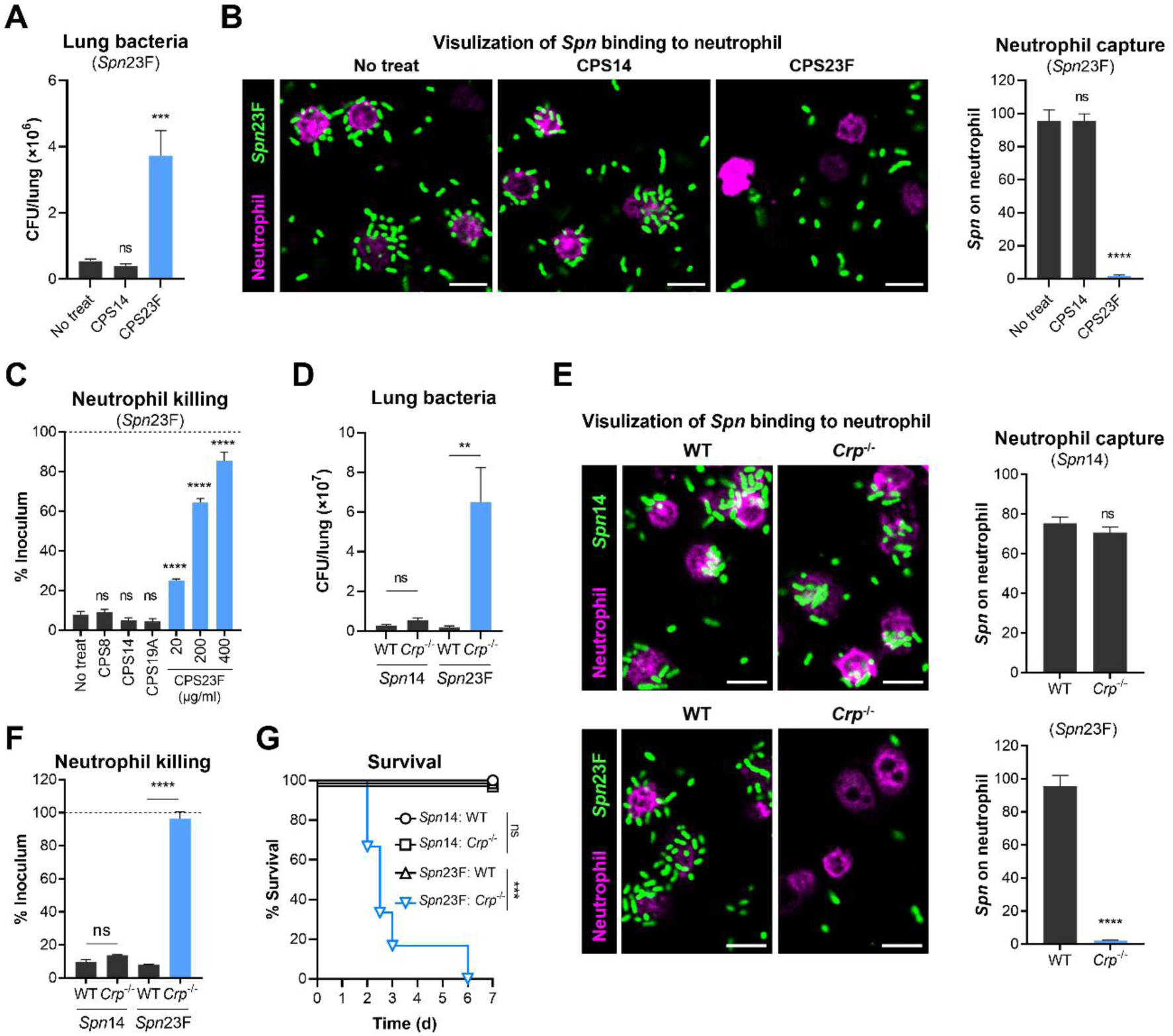
Capsule recognition by plasma CRP for neutrophil immunity to serotype-23F *S. pneumoniae* in the lung. **(A)** Bacterial loads at 12 hr in the lungs under the competitive inhibition of free CPS. Mice were i.n. infected with 10^7^ CFU of *Spn*23F simultaneously with the administration of 150 μg free CPS14 or CPS23F, respectively. n = 6. **(B)** Fluorescent imaging and quantification to show the inhibition of *Spn*23F binding to primary neutrophils by free CPS23F. Free CPS14 or CPS23F was added at a concentration of 400 μg/ml. n = 10 random FOVs. Scale bar, 10 μm. **(C)** Dose-dependent inhibition of *Spn*23F killing by primary neutrophils in the presence of free CPS23F. The heterologous CPS8, CPS14, and CPS19A were used as negative controls and added at a concentration of 400 μg/ml. n = 4-6. **(D)** Bacterial loads at 12 hr in the lungs of WT or *Crp*^-/-^ mice i.n. infected with 5 × 10^7^ CFU of *Spn*14 and *Spn*23F. n = 5. **(E)** Fluorescent imaging and quantification of the *Spn*14 and *Spn*23F binding to primary neutrophils with WT or *Crp*^-/-^ serum. n = 10 random FOVs. Scale bar, 10 μm. **(F)** Killing levels of primary neutrophils to *Spn*14 and *Spn*23F under the conditions of WT or *Crp*^-/-^ serum. n = 4-6. **(G)** Survival time of WT or *Crp*^-/-^ mice i.n. infected with 5 × 10^7^ CFU of *Spn*14 and *Spn*23F. n = 6. Ordinary one-way ANOVA with Dunnett’s multiple comparisons test (A, B, C), unpaired *t* test (D, E, F), log-rank test (G), ns, not significant, **, P < 0.01, ***, P < 0.001, ****, P < 0.0001.

Our recent work has shown that C-reactive protein (CRP) binds to the capsule of serotype-23F *S. pneumoniae*.^13^ CRP is an evolutionarily conserved plasma protein in mammals, which is produced in the liver.^34^ While CRP is known to recognize phosphocholine of teichoic acid (C-polysaccharide) on pneumococcal cell wall,^35,36^ its function has not been characterized in pulmonary defense via the identification of capsules. We thus investigated whether CRP plays a similar role in the pulmonary clearance of *Spn*23F by i.n. infecting CRP knockout (*Crp*^-/-^) mice. The *Crp*^-/-^ mice exhibited a 34-fold increase in lung CFU compared to WT mice at 12 hr. By comparison, *Crp*^-/-^ and WT mice showed similar bacterial levels in the lungs when infected with *Spn*14 (**Fig. 3D**). Given the importance of neutrophils in controlling LV pneumococci in the lung, we further explored the contribution of CRP to the neutrophil-mediated capture and killing of *Spn*23F. In sharp contrast to the potent effect of normal mouse serum in promoting neutrophil capture of *Spn*23F, the serum from *Crp*^-/-^ mice did not show any activity. However, serum from *Crp*^-/-^ and WT mice displayed a comparable activity in enabling neutrophil to capture *Spn*14 (**Fig. 3E**). Accordingly, the CRP-deficiency severely compromised neutrophil killing of *Spn*23F but not *Spn*14 (**Fig. 3F**). All of the *Spn*23F-infected *Crp*^-/-^ mice died in 6 days post i.n. infection with an otherwise non-lethal dose in WT condition, but *Crp*^-/-^ mice survived a parallel infection with the isogenic *Spn*14 strain (**Fig. 3G**). Taken together, these *in vitro* and *in vivo* findings have demonstrated that CRP empowers neutrophils to effectively eliminate serotype-23F pneumococci during lung infection.

### Complement system is necessary for the anti-*Spn*23F immunity of neutrophils

Previous *in vitro* studies have shown that CRP activates the complement system via the classical pathway once binding to the phosphocholine moiety of pneumococcal cell wall.^37,38^ We thus tested if CPS23F-CRP binding initiates the activation of complement system in the lung by using *C3*^-/-^ mice genetically deficient in complement core protein C3 for lung challenge.^39^ *C3*^-/-^ mice carried 10-fold more *Spn*23F bacteria than WT mice in the lung at 12 hpi. In contrast, the lung bacterial loads were comparable between *C3*^-/-^ and WT mice post infection with the HV *Spn*8 (**Fig. 4A**). Under the *in vitro* conditions, *C3*^-/-^ serum did not substantially enhance neutrophil capture of *Spn*23F as compared with WT serum (**Fig. 4B**). Consistently, the lack of C3 resulted in the complete loss of neutrophil killing of *Spn*23F (**Fig. 4C**). This result demonstrated that C3 is essential for the CRP-activated neutrophil clearance of *Spn*23F bacteria.

**Figure 4.**
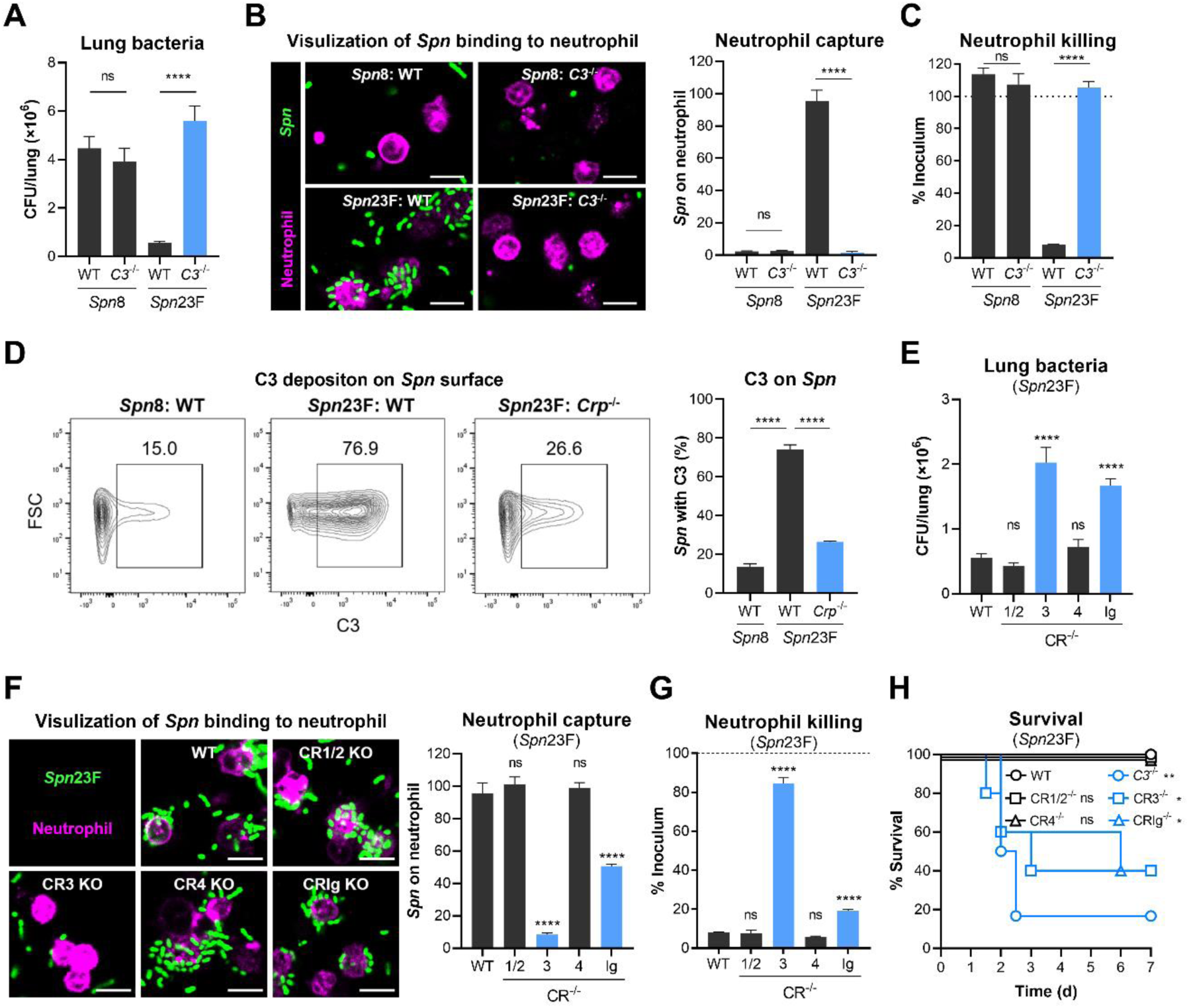
Essential role of complement system for neutrophil-executed *Spn*23F clearance in the lung. **(A)** Bacterial loads at 12 hr in the lungs of WT or *C3*^-/-^ mice i.n. infected with 10^7^ CFU of *Spn*8 and *Spn*23F. n = 5-6. **(B)** Fluorescent imaging and quantification of *Spn*8 and *Spn*23F binding to primary neutrophils with WT or *C3*^-/-^ serum. n = 10 random FOVs. Scale bar, 10 μm. **(C)** Killing levels of primary neutrophils to *Spn*8 and *Spn*23F under the conditions of WT or *C3*^-/-^ serum. n = 4-6. **(D)** Flow cytometry analysis of C3 deposition on *Spn*8 and *Spn*23F surface post incubation with WT or *Crp*^-/-^ serum. n = 3. **(E)** Bacterial loads at 12 hr in the lungs of WT or individual complement receptor KO (CR^-/-^) mice i.n. infected with 10^7^ CFU of *Spn*23F. n = 5-6. **(F)** Fluorescent imaging and quantification of *Spn*23F binding to primary neutrophils isolated from WT or CR^-/-^ mice with the opsonization of WT serum. n = 10 random FOVs. Scale bar, 10 μm. **(G)** Killing levels of primary neutrophils isolated from WT or CR^-/-^ mice to *Spn*23F with the opsonization of WT serum. n = 4-6. **(H)** Survival time of WT, *C3*^-/-^, or CR^-/-^ mice i.n. infected with 5 × 10^7^ CFU of *Spn*23F. n = 5-6. Unpaired *t* test (A, B, C, D), ordinary one-way ANOVA with Dunnett’s multiple comparisons test (E, F, G), log-rank test (H), ns, not significant, *, P < 0.05, **, P < 0.01, ****, P < 0.0001.

C3 promotes immune clearance of microorganisms by depositing on bacterial surface.^40^ Flow cytometry revealed that the presence of normal mouse serum led to more C3 deposition on the surface of *Spn*23F than *Spn*8. However, the lack of CRP resulted in a sharp reduction in C3 molecules on the *Spn*23F surface (**Fig. 4D**). These findings have shown that *Spn*23F is more prone to C3 deposition than *Spn*8 in the presence of CRP.

C3 deposition on bacterial surface is known to enhances the opsonophagocytosis by binding to the complement receptors (CRs) on phagocytes.^40^ Our flow cytometry analysis showed that the lung neutrophils abundantly expressed CR3, followed by CRIg, with low levels of CR1/2 and CR4 (**Fig. S4A**). We evaluated the contribution of CRs to pulmonary immunity against *S. pneumoniae* using mice lacking various CRs. While CR3^-/-^ and CRIg^-/-^ mice showed significant increase in lung bacteria at 12 hpi, mice lacking CR1/2 and CR4 showed similar levels of lung bacteria as WT controls (**Fig. 4E**). This experiment indicated the importance of CR3 and CRIg in lung defense against *Spn*23F. In the *in vitro* neutrophil-bacterial interaction system, neutrophils isolated from the lungs of CR3^-/-^ mice showed almost complete loss of *Spn*23F binding and killing. To a less extent, CRIg^-/-^ mice also displayed an obvious impairment of neutrophil immunity in anti-*Spn*23F ability (**Fig. 4F and 4G**). However, consistent with the marginal expression levels of CR1/2 and CR4 on neutrophils, neutrophils from mice lacking those C3 receptors had comparable levels of *Spn*23F binding and killing as WT mice (**Fig. 4F and 4G**).

Subsequent survival experiments showed that C3, CR3, and CRIg are important for host defense against lung infection of *Spn*23F. Intranasal infection with *Spn*23F led to 83% mortality of the *C3*^-/-^ mice and 60% mortality of CR3^-/-^ and CRIg^-/-^ mice. In contrast, CR1/2- and CR4-deficient mice did not show abnormal susceptibility to *Spn*23F infection (**Fig. 4H**). Together, these findings demonstrated that C3 and its receptors (CR3 and CRIg) functionally link neutrophils and CRP, the binding receptor for serotype-23F capsule, in capturing and killing *Spn*23F bacteria in the lung.

Although the potential plasma receptors specific for capsules of LV *Spn*14 and *Spn*19A are screened in the process, we conducted a preliminary exploration to assess whether the soluble receptors of CPS14 and CPS19A were also involved in the activation of complement system. The primary neutrophils absolutely lost the bactericidal capacity to two LV serotypes (14 and 19A) with C3-deficient serum. Similarly, the CR3- and CRIg-deficient neutrophils showed significantly reduced levels on the killing of *Spn*14 and *Spn*19A (**Fig. S4B and S4C**). These results strongly support that plasma receptors recognize the LV pneumococcal capsules, activate the complement system, and thereby facilitate neutrophil clearance of LV *S. pneumoniae* in the lung.

### Influenza virus enhances the pathogenicity of low-virulence pneumococcal serotypes

It is puzzling that LV pneumococcal serotypes tested in this work are effectively eliminated by neutrophils in the inflammatory phase of lung infection in mice (**Fig. 1**), but these serotypes are highly associated with pneumococcal disease in humans.^41,42^ Although some of the discrepancies could be due to potential host-specific difference in the susceptibility to pneumococcal serotypes, influenza virus co-infection has been well documented to enhance human susceptibility to pneumococcal disease.^18^ We thus tested the hypothesis that the LV pneumococcal serotypes could be converted to more virulent pathogens in mice by influenza virus co-infection, which would explain the phenotypic difference of these serotypes between human and mouse. We tested this possibility by prior i.n. infection of mice with a sublethal dose of mouse-adapted PR8 strain of influenza A virus (IAV) (30 PFU/mouse), and 7 days later with a sublethal dose of isogenic strains of eight LV serotypes (10^7^ CFU/mouse) (**Fig. 5A**). The mice post PR8 single infection exhibited a maximum body weight loss of 10% on day 7, followed by recovery to the original body weight on day 10 (**Fig. S5A**), indicating successful establishment of a mild influenza infection in mice. The virus-infected mice became much more susceptible to secondary bacterial infection of LV pneumococci. All pneumococcal mono-infected mice survived the pulmonary infections with isogenic strains of serotypes 6B, 7F, 9V, 14, 18C, 19A, 19F, and 23F; in contrast, the mice pre-infected with IAV all succumbed to the same challenge within 7 days despite of variations in the mean survival time among the co-infection groups (**Fig. 5B**). CFU enumeration revealed high levels of bacterial burden in the lungs of the co-infected mice post LV pneumococcal infection at 24 hr (**Fig. S5B**). The mean bacterial load increased by approximately 2,000 folds from bacterial mono-infection to viral and bacterial co-infection (**Fig. 5C**). Many of the co-infected mice developed various levels of bacteremia, while bacteria were barely detected in the bloodstream for the mono-infected mice (**Fig. S5C-E**). These results demonstrated that IAV co-infection dramatically enhances the growth and virulence capacity of the LV pneumococcal serotypes during the lung infection.

**Figure 5.**
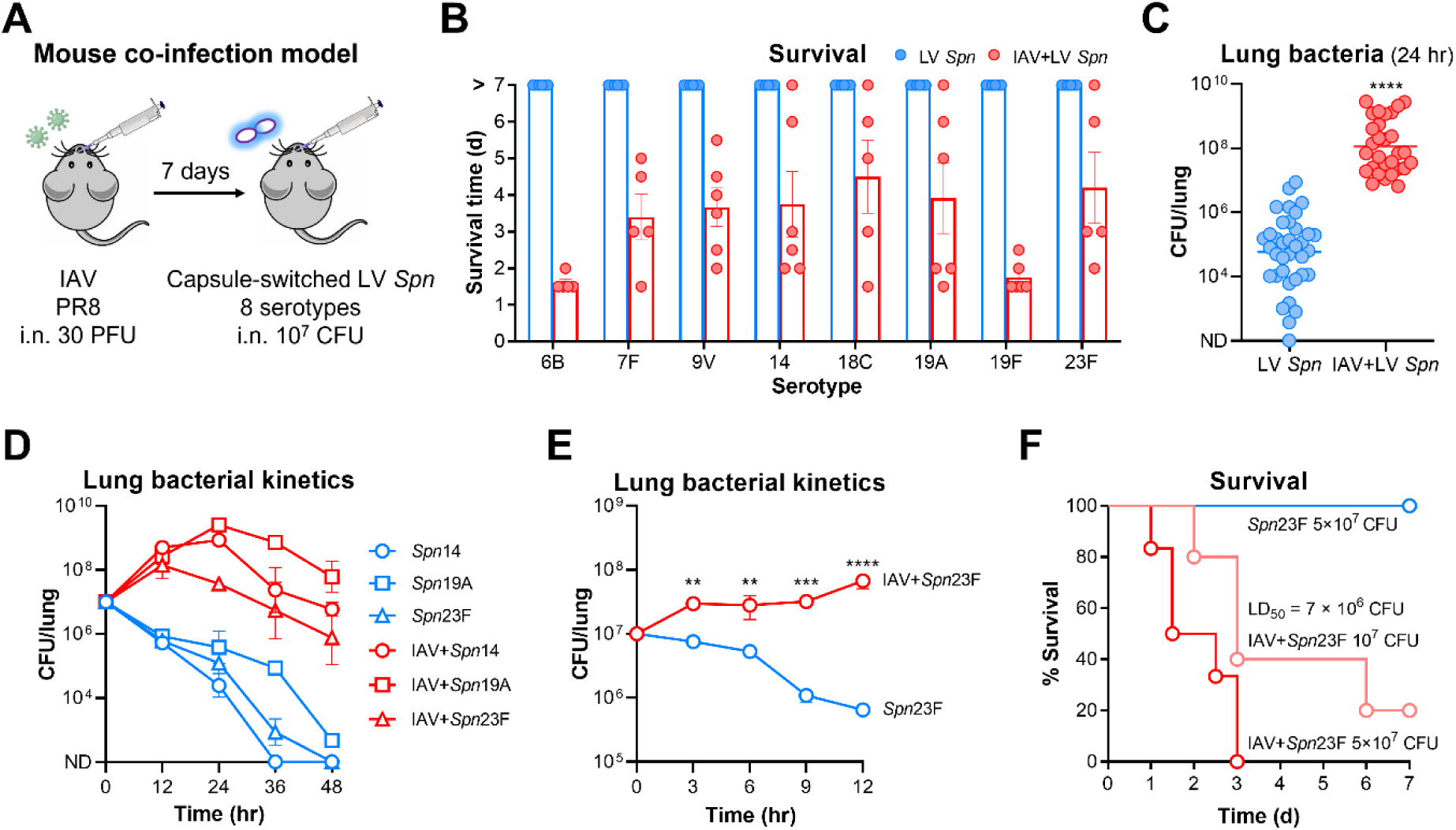
Heightened pathogenicity of LV *S. pneumoniae* serotypes under influenza co-infection. **(A)** Schematic workflow of assessment for the virulence level of capsule-switched LV *S. pneumoniae* strains post influenza A virus (IAV) challenge in mouse co-infection model. **(B)** Survival time of CD1 mice i.n. infected with 10^7^ CFU of LV *S. pneumoniae* strains at 7 days post challenge with 30 PFU of IAV. n = 5-6. **(C)** Bacterial loads at 24 hr in the lungs of mice infected as in (B). Data were combined with 3-6 mice per group for each LV *S. pneumoniae* strain, and each dot represented one mouse. **(D)** Bacterial kinetics within 48 hr in the lungs of mice i.n. infected with 10^7^ CFU of representative LV *Spn*14, *Spn*19A, and *Spn*23F post IAV challenge. n = 3-6. **(E)** Bacterial kinetics within 12 hr in the lungs of mice i.n. infected with 10^7^ CFU of *Spn*23F post IAV challenge. n = 3-6. **(F)** Survival time of mice i.n. infected with 10^7^ or 5 × 10^7^ CFU of *Spn*23F post IAV challenge. LD_50_ during IAV and *Spn*23F co-infection was calculated and shown. n = 5-6. Unpaired *t* test (C), ordinary two-way ANOVA with Sidak’s multiple comparisons test (E), **, P < 0.01, ***, P < 0.001, ****, P < 0.0001.

We further characterized the temporal features of influenza-induced outgrowth of LV pneumococci by monitoring lung bacteria of IAV-primed mice post i.n. infection with representative LV serotypes (14, 19A, and 23F). In sharp contrast to sterilizing immunity in the lungs of mice in the first 48 hpi, the IAV-primed mice showed persistently high levels of pulmonary bacteria (**Fig. 5D**). More detailed sampling in the first 12 hr revealed a striking difference in lung bacterial dynamics between the mono- and co-infected mice. *Spn*23F bacteria in co-infected mice were 4- and 5-fold higher than that in mono-infected mice at 3 and 6 hpi, respectively. This difference became even greater with 29- and 104-fold more bacteria in the lungs of co-infected mice at 9 and 12 hpi (**Fig. 5E**). Further experiments showed that viral pre-infection increased the virulence level of *Spn*23F by 13 folds, in terms of LD_50_ (**Fig. 5F**). These data showed that influenza virus infection dramatically weaken the lung innate immunity against *S. pneumoniae*.

### Influenza virus infection weakens CR3-mediated pneumococcal capture of neutrophils

Because AMs and neutrophils are important in controlling LV serotypes in the onset and inflammatory phases of pneumococcal infection, respectively (**Fig. 2**), we first assessed the number of AMs and lung neutrophils during co-infection. Consistent with the previous study,^43^ IAV-infected mice showed a dramatic loss of AMs in the BALF as compared with the control mice (**Fig. 6A**, 0 hr). The decreased AMs in the virus-infected mice were further reduced following the secondary bacterial infection (**Fig. 6A**, 12 hr). In sharp contrast, neutrophils in the BALF of viral mono-infected mice were strikingly increased (**Fig. 6B**, 0 hr), which were doubled in the inflammatory phase following subsequent bacterial infection (**Fig. 6B**, 12 hr). This drastic increase of neutrophils and simultaneous growth of LV pneumococci in the lungs of co-infected mice strongly suggested that neutrophils in the virus-primed mice are defective in clearing LV *S. pneumoniae* efficiently in the lung. As expected, the *in vitro* bacterial binding assay showed that the lung neutrophils from co-infected mice were significantly impaired in bacterial capture to LV serotypes as compared with those from mice infected with bacteria alone (**Fig. 6C**). This result was supported by significantly lower bacterial killing of lung neutrophils from co-infected mice against LV *S. pneumoniae* strains (**Fig. 6D**). Together, these results demonstrated that bacterial capture immunity of neutrophils is significantly weakened by influenza virus infection.

**Figure 6.**
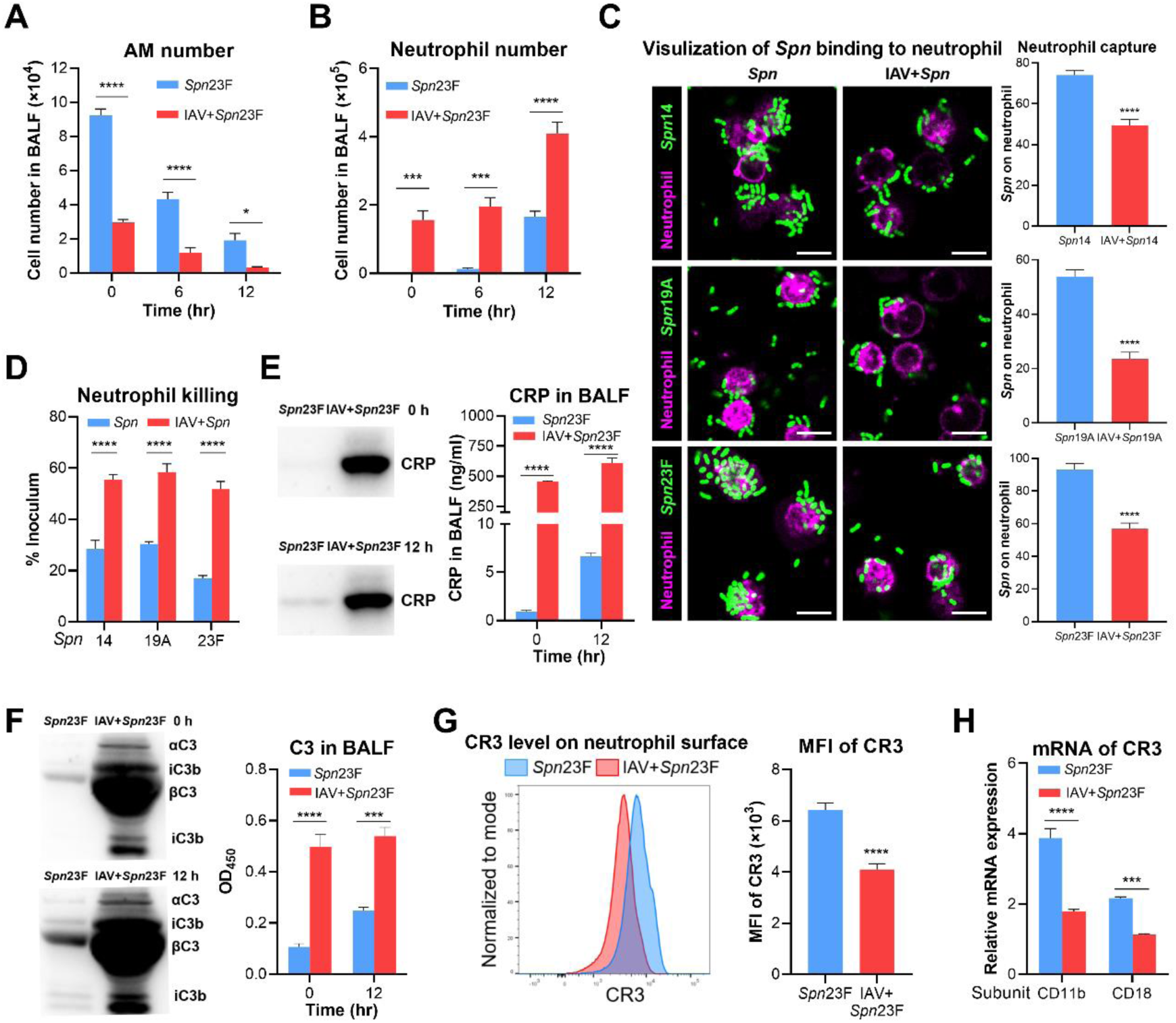
Enhanced immune evasion of LV *S. pneumoniae* serotypes via neutrophil CR3 suppression during influenza co-infection. **(A and B)** AM (A) and neutrophil (B) numbers at 0, 6, and 12 hr in the BALF of C57BL/6 mice i.n. infected with 10^7^ CFU of *Spn*23F or co-infected with 30 PFU of IAV. n = 3. **(C)** Fluorescent imaging and quantification of LV *S. pneumoniae* strains (serotypes 14, 19A, and 23F) binding to primary neutrophils sorted from mono- or co-infected mice. n = 10 random FOVs. Scale bar, 10 μm. **(D)** Killing rate of *S. pneumoniae* strains with LV serotypes by primary neutrophils sorted as in (C). Bacteria were added at an MOI of 0.5. n = 4. **(E)** Immunoblotting (left) and concentration (right) of CRP at 0 and 12 hr in the BALF of mono- or co-infected mice. n = 3. **(F)** Activation (left) and concentration (right) of C3 at 0 and 12 hr in the BALF of mono- or co-infected mice. n = 3. **(G)** Protein level of CR3 on neutrophil surface at 12 hr isolated from the BALF of mono- or co-infected mice. The abundance of CR3 was determined using flow cytometry and presented as median fluorescence intensity (MFI). n = 6. **(H)** Transcriptional levels of CD11b and CD18 subunits of CR3 on neutrophils collected as in (G), and presented as the relative expression to the GADPH gene. n = 4-6. Ordinary two-way ANOVA with Sidak’s multiple comparisons test (A, B, E, F), unpaired *t* test (C, D, G, H), *, P < 0.05, ***, P < 0.001, ****, P < 0.0001.

We next chose to elucidate the mechanisms of influenza-induced outgrowth of serotype- 23F pneumococci based on the abovementioned molecular interactions of *Spn*23F with the host. Given the importance of CRP in mediating neutrophil capture of *Spn*23F, we determined whether influenza virus impacted the CRP level in the lung. In sharp contrast to a barely detectable level of CRP in the lungs of uninfected mice, there was a dramatic increase of CRP in the virus single-infected mice, with approximately 457 ng/ml CRP in the BALF (**Fig. 6E**, 0 hr). At 12 hr post bacterial infection alone, pulmonary CRP level was modestly increased to 6.7 ng/ml. Accordingly, the CRP level was elevated to 611 ng/ml in co-infected mice (**Fig. 6E**, 12 hr). Consistently, immunoblotting results showed that the IAV challenge caused dramatical influx of CRP into the alveoli (**Fig. 6E**). This result showed that it was not the lack of CRP leading to the impaired pneumococcal capture of lung neutrophils in co-infected mice.

Since the complement protein C3 is required for shuffling *Spn*23F bacteria to the C3 receptors on neutrophils, we compared the lung C3 levels between mono- and co-infected mice. Similarly to CRP, both immunoblotting and quantitative evaluation showed that PR8 infection alone resulted in a striking increase of C3 in the BALF samples as compared with uninfected controls (**Fig. 6F**, 0 hr). A similarly high level of C3 was observed in the BALF samples of co-infected mice at 12 hr post bacterial infection, which significantly exceeded the modest induction of C3 by bacterial infection alone (**Fig. 6F**, 12 hr). Likewise, bacteria in the BALF of co-infected mice were more densely coated by C3 than those from mono-infected mice (**Fig. S6A**). These data have ruled out the possibility that a shortage of C3 is the reason for the functional deficiency of lung neutrophils in capturing *Spn*23F bacteria.

Given the crucial roles of CR3 and CRIg in mediating neutrophil capture of CRP/C3- opsonized *Spn*23F, we further compared the levels of these C3 receptors on the lung neutrophils of the co-infected mice by flow cytometry. As compared to the lung neutrophils of *Spn*23F mono-infected mice, the cells of the co-infected mice showed substantially reduced CR3 (**Fig. 6G**). Consistently, the lung neutrophils of co-infected mice showed significantly lower mRNA levels of CD11b and CD18 (**Fig. 6H**), the two subunits of CR3. In contrast, there was a similar level of CRIg between the lung neutrophils of bacterial mono-infected and co-infected mice (**Fig. S6B**), which agreed with subtle changes in CRIg mRNA level following IAV infection (**Fig. S6C**). Taken together, these data strongly suggested that influenza virus enhances the survival of LV *S. pneumoniae* in the lung by downregulating CR3 expression on neutrophils and thereby diminishing the capsule receptor-initiated bacterial capture.

### Pre-viral infection also impairs CRP-driven capture of *H. influenzae* by lung neutrophils

*H. influenzae* is also an encapsulated bacterium commonly associated with the mortality of influenza pneumonia, particularly serotype-b *H. influenzae* (*Hib*).^44^ Our previous study has shown that the *Hib* capsule is recognized by CRP, which drives hepatic clearance of *Hib* from the bloodstream of mice.^13^ We thus determined the contribution of CRP-driven immunity of neutrophils to the *Hib* clearance in the lung, based on the importance of the CRP-neutrophil immune pathway in eliminating the LV pneumococcal serotypes. Depleting neutrophils with 1A8 antibody led to a 5-fold increase in *Hib* bacteria in the lungs at 24 hpi (**Fig. 7A**). In a similar manner, mice lacking CRP, C3 or CR3 also displayed significant impairment in clearing *Hib* in the lungs (**Fig. 7B**), indicating that the CRP-driven bacterial clearance pathway is effective in clearing *Hib* in the lower respiratory tract.

**Figure 7.**
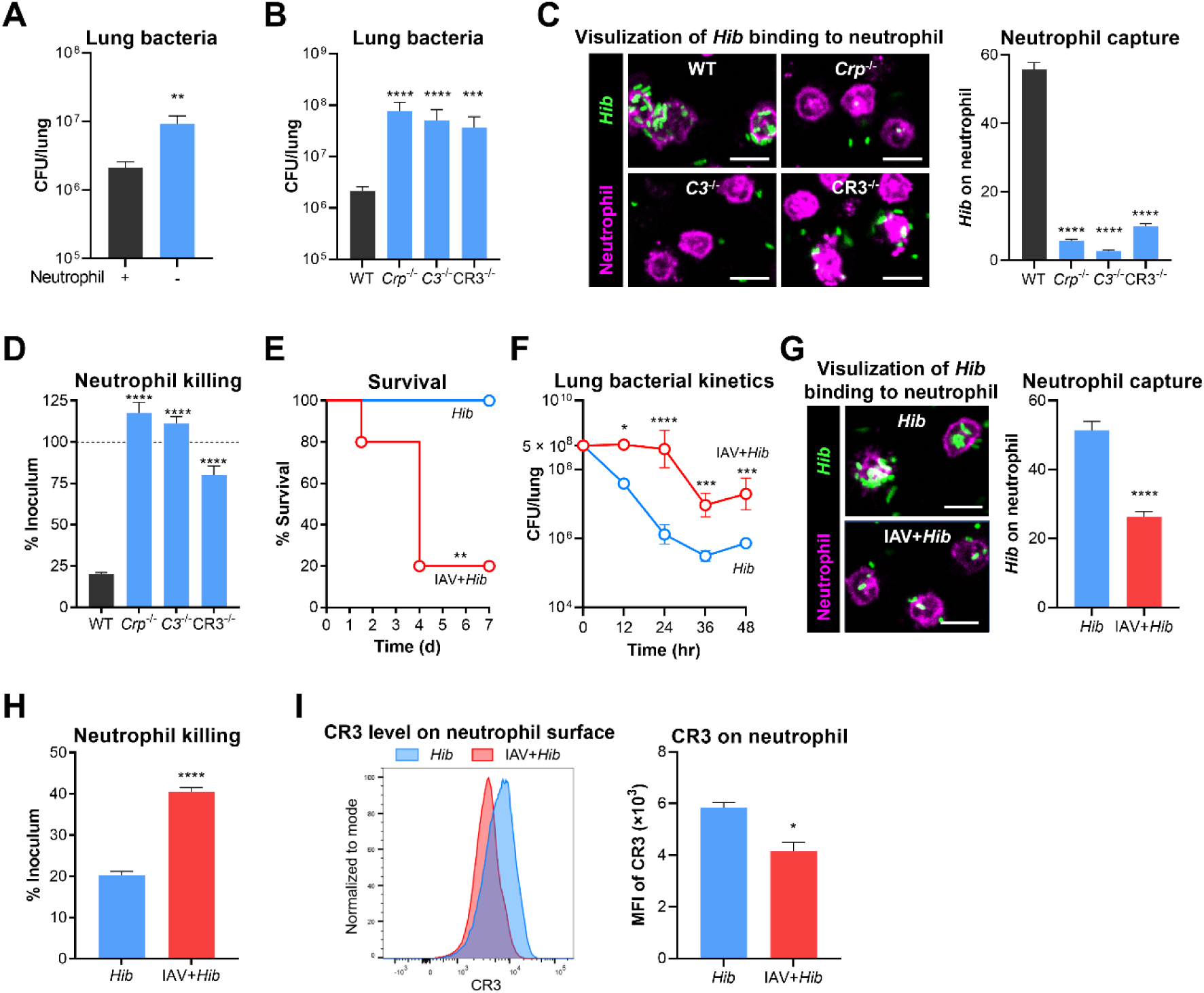
Impairment of CR3-mediated neutrophil clearance of CRP/C3-tagged *Hib* by influenza infection. **(A)** Bacterial loads at 24 hr in the lungs of WT or neutrophil-depleted C57BL/6 mice i.n. infected with 5 × 10^8^ CFU of *Hib*. Neutrophils were depleted by i.p. injection of 500 μg anti-Ly6G antibody (clone 1A8) one day before infection and 100 μg 1A8 antibody at the time of infection. n = 3-6. **(B)** Bacterial loads at 24 hr in the lungs of WT, *Crp*^-/-^, *C3*^-/-^, or CR3^-/-^ mice infected as in (A). n = 5-6. **(C)** Fluorescent imaging and quantification of *Hib* binding to primary neutrophils isolated from WT mice with the addition of WT, *Crp*^-/-^, and *C3*^-/-^ serum, or from CR3^-/-^ mice with the opsonization of WT serum. n = 10 random FOVs. Scale bar, 10 μm. **(D)** Killing rate of *Hib* by primary neutrophils under the conditions as in (C). n = 4. **(E)** Survival time of mice i.n. infected with 5 × 10^8^ CFU of *Hib* at 7 days post challenge with 30 PFU of IAV. n = 5-6. **(F)** Bacterial kinetics within 48 hr in the lungs of mice post infection as in (E). n = 3-6. **(G)** Fluorescent imaging and quantification of *Hib* binding to primary neutrophils sorted from mono- or co-infected mice. n = 10 random FOVs. Scale bar, 10 μm. **(H)** Killing rate of *Hib* by primary neutrophils sorted as in (G). n = 4. **(I)** Protein level of CR3 on neutrophil surface at 12 hr isolated from the BALF of mono- or co-infected mice. n = 3. Unpaired *t* test (A, G, H, I), ordinary one-way ANOVA with Dunnett’s multiple comparisons test (B, C, D), log-rank test (E), ordinary two-way ANOVA with Sidak’s multiple comparisons test (F), ns, not significant, *, P < 0.05, **, P < 0.01, ***, P < 0.001, ****, P < 0.0001.

The functions of CRP-driven anti-*Hib* immunity were further verified through an *in vitro* pathogen-host interaction approach. Lung neutrophils were highly capable of catching *Hib* in the presence of normal mouse serum (**Fig. 7C**). However, the sera from *Crp*^-/-^ or *C3*^-/-^ mice completely failed to drive *Hib* bacteria to neutrophils. In a similar fashion, neutrophils from CR3^-/-^ mice showed dramatic impairment in bacterial capture in the presence of normal serum (**Fig. 7C**). Accordingly, the genetic deficiency in CRP, C3, or CR3 also resulted in profound loss of neutrophil-mediated bacterial killing (**Fig. 7D**). Consistent with the activity of CRP in activating C3 to promote bacterial capture of neutrophils, the flow cytometry analysis revealed significant reduction of C3 deposition on *Hib* cells that were incubated with *Crp*^-/-^ serum as compared with normal serum (**Fig. S7A**). In the context of the CRP-driven neutrophil immunity against serotype-23 *S. pneumoniae* in the inflammatory phase of lung infection (**Fig. 3**), these findings demonstrated that the same immune pathway broadly operates against multiple invading encapsulated bacteria in the lower airway.

We finally determined the impact of influenza virus infection on the capsule receptor-mediated neutrophil immunity against *Hib* in the lung. Pre-infection with PR8 significantly enhanced the *Hib* virulence in co-infected mice. With a non-lethal infection dose of *Hib* (5 × 10^8^ CFU) in normal mice, 80% of the PR8-primed mice succumbed to the secondary bacterial infection (**Fig. 7E**). CFU enumeration showed that *Hib* bacteria were gradually cleared from the lung during the mono-infection; by comparison, there were significantly higher levels of bacterial burden in the lungs of co-infected mice (**Fig. 7F**). Moreover, mice showed marginal levels of *Hib* bacteremia in co-infected mice (**Fig. S7B**), despite heavy bacterial burden in the lungs, indicating that the influenza-compromised immunity against *Hib* mainly operates in the lower respiratory tract.

We further verified whether impaired capacity of lung neutrophils in *Hib* capture is responsible for higher bacterial burden in the lungs of co-infected mice. As compared with the immune cells from *Hib* mono-infected mice, the lung neutrophils from co-infected mice showed significantly reduced level of bacterial binding (**Fig. 7G**) and killing (**Fig. 7H**). The flow cytometry analysis revealed a significant decrease of the CR3 protein on the cell surface of neutrophils isolated from the co-infected mice at 24 hpi (**Fig. 7I**). This result indicated that influenza-induced CR3 repression also accounts for the enhanced susceptibility of mice to secondary infection by *Hib*. Taken together, these data verified that the CRP-driven capsule recognition enables neutrophils to capture and eliminate both Gram-positive (*S. pneumoniae*) and -negative (*H. influenzae*) pathogens in the lungs during the inflammatory phase of infection; furthermore, this anti-bacterial defense pathway is significantly compromised by pre-infection of influenza virus.

## DISCUSSION

Certain serotypes of encapsulated bacteria are more associated with severe pneumonia than other serotypes,^6,45^ but the mechanism behind the association between capsule serotype and virulence in the pathogenesis of bacterial pneumonia are largely undefined. This study has uncovered that capsule serotype determines the virulence level of respiratory bacteria in mouse pneumonia model using isogenic capsule-switched strains. While the high-virulence serotypes are able to persist and grow in the lungs, the low-virulence counterparts are effectively cleared. We further found that infiltrating neutrophils are the major cellular determinant of capsule type-dependent differences in virulence. At the molecular level, capsule-specific receptors via complement activation determine the neutrophil-mediated bacterial capture. The importance of lung neutrophils in the clearance of the low-virulence serotypes is manifested by the hyper-susceptibility of influenza-primed mice to the secondary bacterial infections. This study has not only clarified the mysterious mechanism behind the serotype-dependent virulence dichotomy of encapsulated bacteria in lung infection, but has also shed novel light on the well-publicized synergism between influenza virus and respiratory bacteria.

### Capsule types define pneumococcal virulence in lung infection

A large body of literature has documented a strong association of capsule types with pneumococcal virulence in severe bacterial pneumonia.^8,46,47^ However, it remains unclear how structural variations of capsular polysaccharides mechanistically impact bacterial virulence during lung infection. This work unequivocally demonstrated that capsule types dictate the virulence phenotypes of *S. pneumoniae* using isogenic strains producing various serotypes. While the LV serotypes were gradually cleared from the lungs of mice, the HV serotype strains persisted in the lungs, which led to bacterial outgrowth. Thirteen serotypes tested in the lung infection model showed the same virulence phenotypes as previously observed in the liver.^10^ As demonstrated with the common role of CRP in driving the capture of serotype-23F *S. pneumoniae* by neutrophils in the lung (this work) and by KCs in the liver,^10^ this common phenotypes of LV pneumococcal serotypes in two host niches strongly suggests that the same capsule-binding receptors drive host immunity in both the lung and liver.

### Lung neutrophil is the major immune cell that shapes the virulence phenotypes of *S. pneumoniae*

While neutrophils are recruited to the lungs in response to *S. pneumoniae* infection,^28^ their precise contribution to bacterial clearance is not fully clear. Some studies show that neutrophils are important for host resistance to *S. pneumoniae*,^31,48,49^ but other works have reported that the immune cells have marginal impact on bacterial viability and host survival.^50–52^ This work has uncovered that neutrophil immunity against *S. pneumoniae* in the lungs varies highly, depending on capsule types/structures. Using serotype 23F strain and the corresponding capsule receptor CRP, we have demonstrated that lung neutrophils have poor affinity to capsular polysaccharides and cannot effectively capture encapsulated bacteria unless the bacteria are first recognized by the serotype-specific capsule receptor (see below). Likewise, the HV serotypes that lack specific receptors successfully escape neutrophil capture and phagocytic killing. Our finding has provided a molecular explanation for the prevalence of certain *S. pneumoniae* serotypes in severe pneumonia cases.^7,8,53^ Furthermore, this serotype-dependent phenomenon can also explain the discrepancies regarding the neutrophil immunity among different studies. The past investigations have indifferently used various serotypes of *S. pneumoniae* to study host-pathogen interactions in lung infections,^31,48–52^ which would yield different data, depending on the virulence phenotypes of particular serotypes.

### Capsule-binding receptors are the major molecular determinants of *S. pneumoniae* virulence levels in lung infection

A number of ligand-binding receptors have been identified for macrophage binding to microorganisms, the first step of phagocytosis,^54^ but the pathogen-binding receptors that initiate neutrophil phagocytosis of pathogens are much less understood. Fc receptors promote neutrophil uptake of bacteria bound by IgA or IgG.^55,56^ CR3 is reported to enhance human neutrophil binding to and uptake of *Streptococcus pyogenes* in the presence of nonimmune serum,^57^ but the mechanism of the CR3-mediated bacterial phagocytosis remains to be defined. This work showed that lung neutrophils in general have poor affinity to capsular polysaccharides, and therefore are unable to directly capture encapsulated pneumococci for phagocytic killing. Instead, as demonstrated by the CRP-mediated neutrophil capture of serotype-23 *S. pneumoniae* and serotype-b *H. influenzae*, neutrophils rely on plasma capsule-binding receptors to recognize bacterial capsules to initiate pathogen capture. Similar to its role in initiating pneumococcal capture by KCs,^13^ plasma CRP indirectly promotes neutrophil capture of encapsulated pathogens by activating C3 on bacterial surface and thereby shuffling C3-opsonized bacteria to the C3 receptors. Phenotypically, the intrinsic capsule-binding receptors mainly determine the fates and virulence levels of encapsulated bacteria. The capsule types with intrinsic receptors are translated into the LV phenotypes, while those without specific receptors successfully escape neutrophil capture and possess the HV phenotypes. Although this work has mainly focused on CRP-binding interactions with capsular polysaccharides of serotype-23 *S. pneumoniae* and serotype-b *H. influenzae*, the neutrophil capture-dependent virulence in bacterial lung infections may be applicable to other encapsulated bacteria, because the capsule receptor-mediated immunity has explained the low-virulence phenotypes of many other LV capsule types of encapsulated bacteria in septic infection models.^10,12,13^

### Complement system functionally links capsule-binding receptors and neutrophils

The complement system, composed of C3 and many related proteins, is activated by the classical, alternative, or lectin pathway.^58^ While humans with genetic deficiencies in the complement activation are hyper-susceptible to infections of *S. pneumoniae*, *H. influenzae, Neisseria meningitidis*, and other encapsulated bacteria,^59^ the precise mechanisms of the complement’s anti-infection actions are not fully understood. The alternative pathway is reported to play an important role in innate immunity against *S. pneumoniae*,^60^ whereas the classical pathway is later shown to be more dominant.^61^ A previous study also shows that certain serotypes of *S. pneumoniae* (e.g., serotypes 1, 3, 4, and 8) are more resistant to C3 deposition than others (e.g., serotypes 7, 12, 14, and 25).^62^ Our data have demonstrated that the potency of the complement system against *S. pneumoniae* in lung infection cannot be generalized since it varies greatly, depending on capsule types. Consistent with the previous observation with human C3,^62^ mouse C3 is highly activated on the surface of low-virulence (serotype 23) but not high-virulence (serotype 8) pneumococci. Using CRP as a model, we further uncovered that C3 activation on *S. pneumoniae* and *H. influenzae* is only initiated by the prior binding of capsule-binding receptors to capsular polysaccharides. C3-opsonized bacteria are in turn captured by C3 receptors on neutrophils, leading to phagocytic clearance. Likewise, the lack of specific receptors to the capsules of the high-virulence serotypes is responsible for their poor opsonization by C3 and thereby evasion of neutrophil’s phagocytic clearance. In short, the complement system fulfills its immune function against low-virulence encapsulated bacteria by acting as a liaison between capsule-binding receptors and C3 receptors on neutrophils. Our finding provides an explanation for the discrepancies in the current literature regarding the specific contributions of the complement system to various serotypes of encapsulated bacteria.

### Alveolar macrophages serve as house-keeping phagocytes against the HV and LV encapsulated bacteria in the lower airway

AMs are vital for the clearance of pulmonary bacteria,^63^ but capsules significantly inhibit AM’s phagocytosis of encapsulated bacteria.^64^ The precise impact of capsular structural variation on the anti-bacterial activity of AMs remains undefined. This study shows that AMs alone are able to eliminate both the high- and low-virulence serotypes of *S. pneumoniae* at a similar pace in the early phase of lung infection. Both the HV and LV pneumococci were kept to a similarly steady level in the lungs of normal mice in the first 6 hr post intranasal infection when AMs represented the vast majority of phagocytes in the lower airway; two bacteria outgrew similarly in the lungs of mice lacking AMs during the same period. Consistent with the apoptosis-driven bactericidal mechanism of AMs,^65–67^ the cellular number of AMs substantially but similarly declined in the lungs of mice infected with either HV or LV pneumococci in the early phase of infection. The gradual loss of AMs coincided with the outgrowth of the HV serotypes in the lungs in the inflammatory phase of infection (post 6 hr). During the same later infection period, rapid infiltration of neutrophils led to gradual disappearance of the LV pneumococci from the lungs. The different dynamics and bactericidal activities of AM and neutrophils during lung infection can explain the phenotypic variations among pneumococcal serotypes.

The slow pace of AM’s bactericidal activity is reminiscent of what has been observed with the splenic red pulp macrophage,^68^ which is in sharp contrast with rapid bacterial clearance of KCs.^10,11^ Similar to the receptor-mediated capsule binding of neutrophils shown in this work, KCs bind to the capsules of the LV pneumococcal serotypes via various capsule-binding receptors.^10,12,13^ In contrast, red pulp macrophages cannot bind to capsules and instead capture pneumococci by natural antibody-mediated recognition of capsule-shielded cell wall phosphocholine.^68^ These lines of information suggests that AMs are unable to specifically recognize capsular polysaccharides for phagocytic clearance of encapsulated bacteria. The precise mechanism of the AM’s phagocytic action toward encapsulated bacteria warrants further investigations.

### Influenza virus-induced neutrophil dysfunction enhances the survival and pathogenicity of low-virulence bacteria in the lung

The virus-induced hyper-susceptibility of secondary bacterial infections has been previously attributed to impaired functions of AMs and epithelial cells.^21–23^ This study has added the important contribution of virus-induced neutrophil dysfunction to this viral-bacterial synergism in pathogenesis. While this virus-induced immunosuppression may have a general effect on the complement-mediated phagocytic activity of neutrophils, our data showed that it has the most damaging impact on the capsule receptor-driven capture of the low-virulence encapsulated bacteria. In contrast to the potent clearance of the low-virulence pneumococcal serotypes by lung neutrophils of normal mice, the neutrophils from the lungs of virus-primed mice had significantly reduced capacity in capturing C3-opsonized bacteria, due to the virus-induced disappearance of CR3, the major C3 receptor on lung neutrophils. This finding may explain the serotype shift of *S. pneumoniae* isolated from the lobar pneumonia patient during the 1918 influenza pandemic. Prior to the pandemic, serotypes 1, 2, and 3 were the dominant isolates,^7^ but the other serotypes were much more frequently found among patients with pneumococcal pneumonia during the influenza pandemic.^9^

### Limitations of this study

While our study has provided significant insights into the molecular and cellular mechanisms of bacterial pneumonia pathogenesis, it has several limitations. The mouse infection models may not fully reflect pathogen-host interactions and disease outcomes in humans. In addition, the *S. pneumoniae* and *H. influenzae* serotypes and host receptor (CRP) studied in this work may not completely reflect the behaviors of the remaining serotypes and the corresponding receptors. These limitations will be addressed by using humanized models and identifying capsule-binding receptors for additional serotypes.

## STAR METHODS

### RESOURCE AVAILABILITY

#### Lead contact

Additional information and requests including materials and resources are available by contacting the lead contact, Jing-Ren Zhang.

#### Materials availability

All materials are available upon request, including bacterial and mouse strains generated in this work.

### EXPERIMENTAL MODEL DETAILS

#### Mice

All infection experiments were conducted in 6-8 weeks old female C57BL/6 or CD1 mice purchased from Charles River (Beijing, China) following the animal protocols of the Institutional Animal Care and Use Committee in Tsinghua University. All knockout mice were maintained as homozygous lines on the C57BL/6 background. *C3*^-/-^ mice were purchased from the Jackson Laboratory (Maine, USA). *Crp*^-/-^ and CR1/2^-/-^ (*Cd21*^-/-^) mice were obtained from the Gempharmatech (Nanjing, China). CRIg^-/-^ (*Vsig4*^-/-^) mice were acquired from Genentech (San Francisco, USA). CR3^-/-^ (*Itgam*^-/-^), CR4^-/-^ (*Itgax*^-/-^), and AM-cKO (*Cd11c*-Cre*Hdac1*^fl/fl^*Hdac2*^fl/fl^) mice were generated in our previous studies.^33,69^ All mice were housed under specific pathogen-free conditions.

#### Bacteria and viruses

All bacterial and viral strains of *S. pneumoniae*, *H. influenzae*, and influenza A virus used in this study are described in **Table S1**.

#### Cell lines

Madin-Darby canine kidney (MDCK) cells were maintained at 37 °C with 5% CO_2_ in DMEM medium (Gibco) complemented with 10% FBS (Viva Cell Biosciences), 1% GlutaMAX (Gibco), and 1% penicillin/streptomycin (Gibco), as previously described.^21^

### METHOD DETAILS

#### Bacterial and viral cultivation

*S. pneumoniae* strains were cultured in Todd-Hewitt broth (Oxoid) with 0.5% yeast extract (Oxoid) or tryptic soy agar plates (BD Difco) with 3% defibrinated sheep blood as depicted previously.^10^ *H. influenzae* strain was incubated in brain-heart infusion (BHI) broth (BD Difco) or on BHI agar with 1.5% agar supplemented with 10 µg/ml hemin (Sigma-Aldrich) and 1 µg/ml NAD (Sigma-Aldrich) as described.^70^ Influenza A virus strain was grown in 9-day embryonated hen’s eggs and titrated by plaque assays on MDCK cells as described elsewhere.^21^

#### Mouse infection

Lung infections were carried out by intranasal (i.n.) inhalation with 30 μl of PBS after anesthesia as described previously.^21^ Bacteria were prepared using frozen stocks with pre-determined viable bacterial concentrations. CPS blocking assay was conducted by i.n. inhalation with 150 μg of free CPS together with bacterial inoculum. The co-infection procedure was performed as follows: mice were i.n. infected with 30 PFU of influenza virus; 7 days later with 10^7^ CFU of *S. pneumoniae*, or 5 × 10^8^ CFU of *H. influenzae*.

Bacteria in the blood were determined by retroorbital bleeding and CFU plating. Bacterial loads in the lungs were estimated by sacrificing mice at indicated time points to prepare tissue homogenates. Viable bacteria were enumerated by plating serial dilutions of each sample onto blood agar plates. Survival rates were monitored in 7 days post infection or mice reached a humane endpoint (body weight loss >25%). LD_50_ values were determined with the combinations of infection dose and survival rate using an LD_50_ calculator (AAT Bioquest).

#### Immune cell depletion

Alveolar macrophages (AMs) were depleted by i.n. inhalation with 50 μl of clodronate liposome (LIPOSOMA) 5 days before pneumococcal infection as reported.^71^ Neutrophils were depleted by intraperitoneal (i.p.) injection of 500 μg anti-Ly6G antibody (clone 1A8, BioXCell) one day before infection and daily injection of 100 μg 1A8 antibody until the end of the experiments as described.^10^ Depletion efficiency of AMs and neutrophils in the lungs was more than 90% as verified by flow cytometry.

#### Immune cell analysis by flow cytometry

Immune cells in the lungs of mice were analyzed by flow cytometry as described.^71^ Cells in the alveolar space of mice were assessed by lavaging the lung six times with 600 μl phosphate-buffered saline (PBS) to collect bronchoalveolar lavage fluid (BALF). Cells in BALF were enriched by centrifugation at 500 g and lysis of red blood cells using RBC lysis buffer (BioLegend) on the ice for 1 min. IgG receptors on cells (per 5 × 10^5^) were blocked with 1% anti-CD16/32 antibody (BioLegend) in 25 μl FACS buffer (PBS supplemented with 3% FBS) for 10 min, and then treated with additional 0.25 μl cell typing antibodies in 25 μl FACS buffer for 20 min: APC-Cy7 anti-CD45, PE anti-SiglecF, FITC anti-Ly6G, eFluor 450 anti-Ly6C, BV605 anti-CD11b, and PE-Cy7 anti-CD11c. The cells were gated as AMs (CD45^+^SiglecF^+^CD11c^+^), neutrophils (CD45^+^Ly6G^+^Ly6C^+^CD11b^+^), and monocytes (CD45^+^Ly6G^-^Ly6C^high^CD11b^+^). Total leukocyte numbers were enumerated using a hemocytometer.

#### Bacterial fluorescence labeling

Bacterial strains were labeled with fluorescence of FITC or AF647 as previously described.^10^ Briefly, 10^8^ CFU of bacteria were collected and resuspended in 100 μl PBS with the supplement of 200 μg/ml FITC or AF647 (Sigma-Aldrich). The complex was incubated in the dark for 30 min. The free fluorescence was removed by washing three times with 1 ml PBS. Higher than 90% efficiency of fluorescence labeling on bacteria was confirmed by flow cytometry.

#### Assessment of *in vivo* bacterial capture by immune cells

Bacterial capture by immune cells in the lungs was evaluated *in vivo*, modified from a previous study.^72^ Following the infection of *S. pneumoniae* labeled with AF647 fluorescence or expressing RFP protein at indicated time points, immune cells from the BALF samples were collected and analyzed by flow cytometry to quantify those with or without bacterial fluorescence.

#### Bacterial binding and killing by lung neutrophils

Bacterial binding to neutrophils was carried out principally as described.^71,72^ Briefly, neutrophils were isolated from the lungs of mice infected with *Spn*23F at 12 hpi by passing shredded lung tissues through a 70-μm cell strainer to obtain single cells. The cells were enriched by centrifugation at 500 g for 5 min, lysis of red blood cells with 1 ml RBC lysis buffer on the ice for 3 min, and incubation in 25 μl FACS buffer (per 5 × 10^5^ cells) containing 1% anti-CD16/32 antibody for 10 min to block IgG receptors. Neutrophils were labeled by adding 0.25 μl AF647 anti-Ly6G (BioLegend) in 25 μl FACS buffer and incubating for 20 min. Cells were pelleted, resuspended in RPMI 1640 medium (Corning) without FBS.

For bacterial binding, the labeled cells were seeded into a 4-chamber glass bottom dish (Cellvis) at 3 × 10^5^ cells/well to enrich adherent neutrophils at 37 °C with 5% CO_2_ for 30 min. After removing non-adhering cells by washing three times with RPMI 1640 medium, about 80% of the remaining adherent cells were Ly6G^+^ neutrophils with polymorphonuclear morphology, as confirmed by fluorescence microscopy (**Fig. S2F and S2G**). FITC-labeled bacteria were added to adherent cells in a multiplicity of infection (MOI, bacterium vs cell) of 10 in the presence or absence of 500 μl BALF supernatant or 20% mouse serum. The plate was centrifuged at 500 g for 5 min to maximize binding interactions between neutrophils and bacteria, incubated for 30 min at 37 °C with 5% CO_2_, and observed by confocal microscopy. Bacterial killing by neutrophils was assessed in a similar manner, except for seeding 5 × 10^5^ cells into a 48-well culture plate in the initial step, using an MOI of 0.2 in the infection step and quantifying free (in the supernatant) and adherent (on the cells) bacteria by CFU plating at the last step. Bacterial killing rate was calculated by dividing the remaining (free and adherent) viable bacteria by the inoculum (∼ 10^5^ CFU).

Considering the complexed cellular population after IAV infection, bacterial binding and killing of lung neutrophils from mice infected with *Spn*23F alone or the IAV-*Spn*23F combination were assessed with neutrophils isolated by FACS sorting of Ly6G^+^ cells when compared between mono- and co-infection groups. The purity of sorted neutrophils was more than 95% as confirmed by flow cytometry.

#### Detection of CRP and C3 in BALF

The relative content of CRP and C3 in BALF samples was determined using western blotting according to the previous studies.^73,74^ Briefly, BALF supernatants were mixed with SDS-PAGE loading buffer, boiled for 10 min, and then loaded into the SDS-PAGE gel. Gels were transferred to polyvinylidene difluoride (PVDF) membrane (Bio-Rad). CRP was detected by rabbit anti-mouse CRP IgG antibody (Invitrogen) and HRP-conjugated goat anti-rabbit IgG (Easybio); C3 by HRP-conjugated goat anti-mouse C3 IgG antibody (Cappel). The quantification of CRP in BALF samples was determined by commercial enzyme-linked immunosorbent assay (ELISA) kit (R&D Systems) following the manufacturer’s instructions. The concentration of C3 was measured based on the previous study.^75^

#### C3 deposition on bacteria

C3 deposition on bacterial cells was detected as described.^73,74^ In brief, bacteria (5 × 10^7^ CFU) were incubated with 10% WT or *Crp*^-/-^ serum in 500 μl RPMI 1640 medium at 37 ℃ for 30 min, and washed with PBS to remove unbound C3. Bacterial pellets were suspended with 100 μl PBS containing 1,000-fold diluted goat anti-mouse C3 IgG antibody (Cappel) and incubated for 20 min, followed by treating with 100 μl PBS containing 1,000-fold diluted AF568 donkey anti-goat IgG antibody (Invitrogen). The level of C3 deposition on bacteria was detected by flow cytometry.

#### Quantification of C3 receptor mRNA

The mRNA levels of C3 receptors were analyzed by quantitative reverse transcription polymerase chain reaction (qRT-PCR) as described.^76^ The total RNA was extracted from mouse neutrophils by TRIzol reagent (Invitrogen) following the manufacturer’s instructions. cDNAs were obtained with the neutrophil RNA samples using HiFiScript cDNA Synthesis Kit (CWbiotech). The following quantitative assessment was conducted with SYBR Green Mixture (CWbiotech) and C3 receptor-specific primers. Relative gene expressions of C3 receptors were normalized with the expression of GADPH. All the primers used for the qRT-PCR are listed in **Table S2.**

### QUANTIFICATION AND STATISTICAL ANALYSIS

Statistical analysis was carried out using GraphPad Prism software (version 9.0). All values are presented as Mean ± SEM. Unpaired *t* test was utilized to compare the statistical significance between two groups. Ordinary one-way ANOVA with Dunnett’s multiple comparisons test or two-way ANOVA with Sidak’s multiple comparisons test were performed to analyze multiple groups. Statistical tests, sample numbers and P values for each experiment are noted in the figure legend.

## ACKNOWLEDGEMENTS

We thank the Tsinghua research platforms for assistance in animal experimentation (Laboratory Animal Research Center), flow cytometry (Center for Cell Biology), and IVM imaging (Center for Cell Biology). Graphical abstract was created in BioRender (Fang, Y., 2024, https://BioRender.com/q93o668). This work was supported by grants from the National Key R&D Program of China to J.-R.Z. and H.A. (2023YFC2306300 and 2023YFC2308003), the National Natural Science Foundation of China to J.-R.Z. (82330071), and the Peking University Medicine Sailing Program for Young Scholars’ Scientific & Technological Innovation and the Fundamental Research Funds for the Central Universities to H.A. (BMU2024YFJHPY011).

## AUTHOR CONTRIBUTIONS

Conceptualization: Y.F., H.A., J.-R.Z.; experimentation: Y.F., W.M., C.Q., J.H., X.T.; methodology: Y. F., X.H., W.L., L.W., H.A.; data analysis: Y. F., W.M., H.A.; writing original draft: Y.F., W.M., H.A., J.-R.Z.; supervision: J.-R.Z.; funding: H.A. and J.-R.Z.

## SUPPLEMENTAL FIGURE LEGENDS

**Figure S1.**
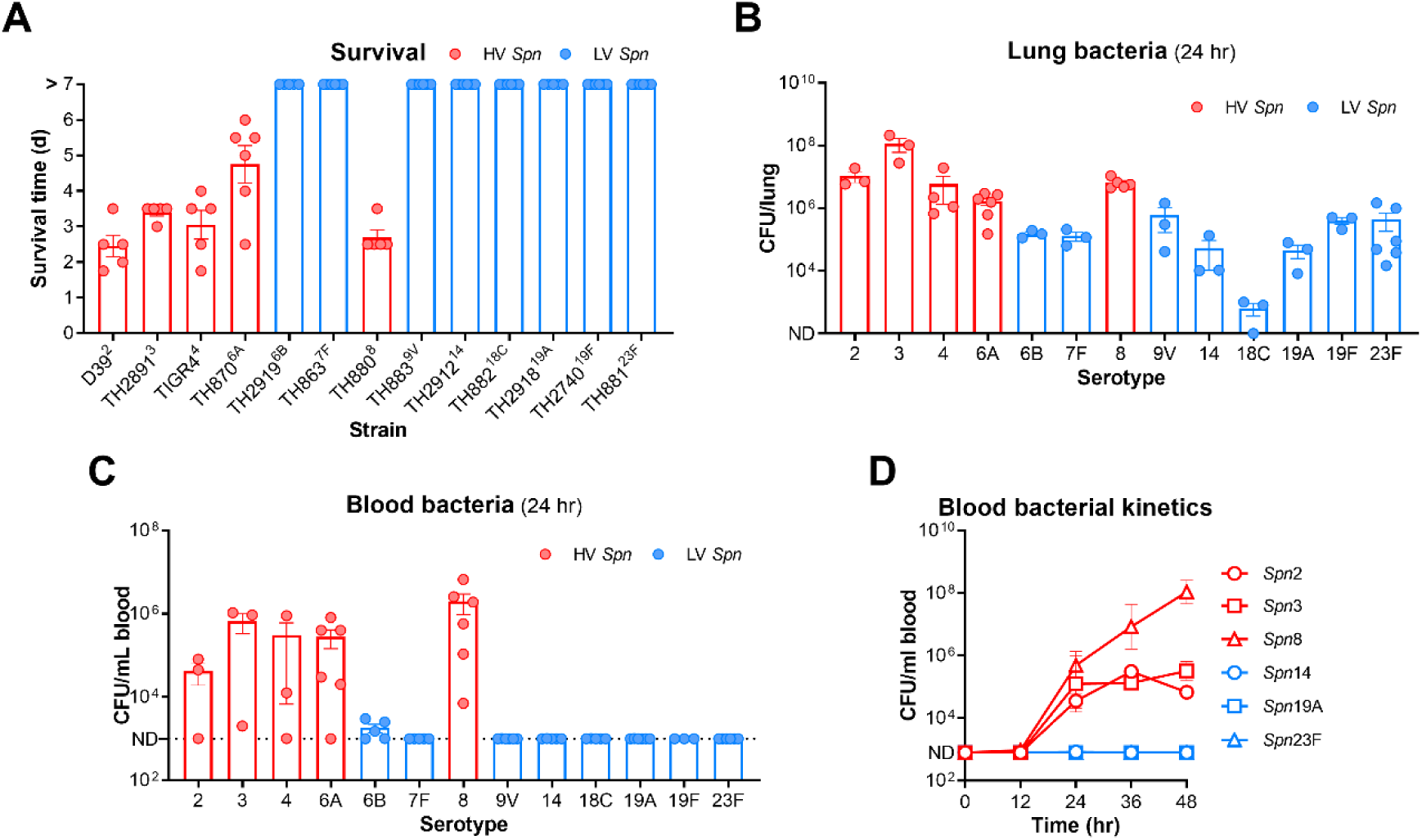
Capsule type-dependent pneumococcal virulence in lung infection. **(A)** Survival time of CD1 mice i.n. infected with 10^7^ CFU of wild-type *S. pneumoniae* strains. HV and LV serotypes were indicated with red and blue dots, respectively. n = 5-6. **(B and C)** Bacterial loads at 24 hr in the lungs (B) and blood (C) of mice infected as in (A). n = 3-6. **(D)** Bacterial kinetics within 48 hr in the blood of mice i.n. infected with 10^7^ CFU of HV (serotypes 2, 3, and 8) and LV (serotypes 14, 19A, and 23F) strains. n = 3-6.

**Figure S2.**
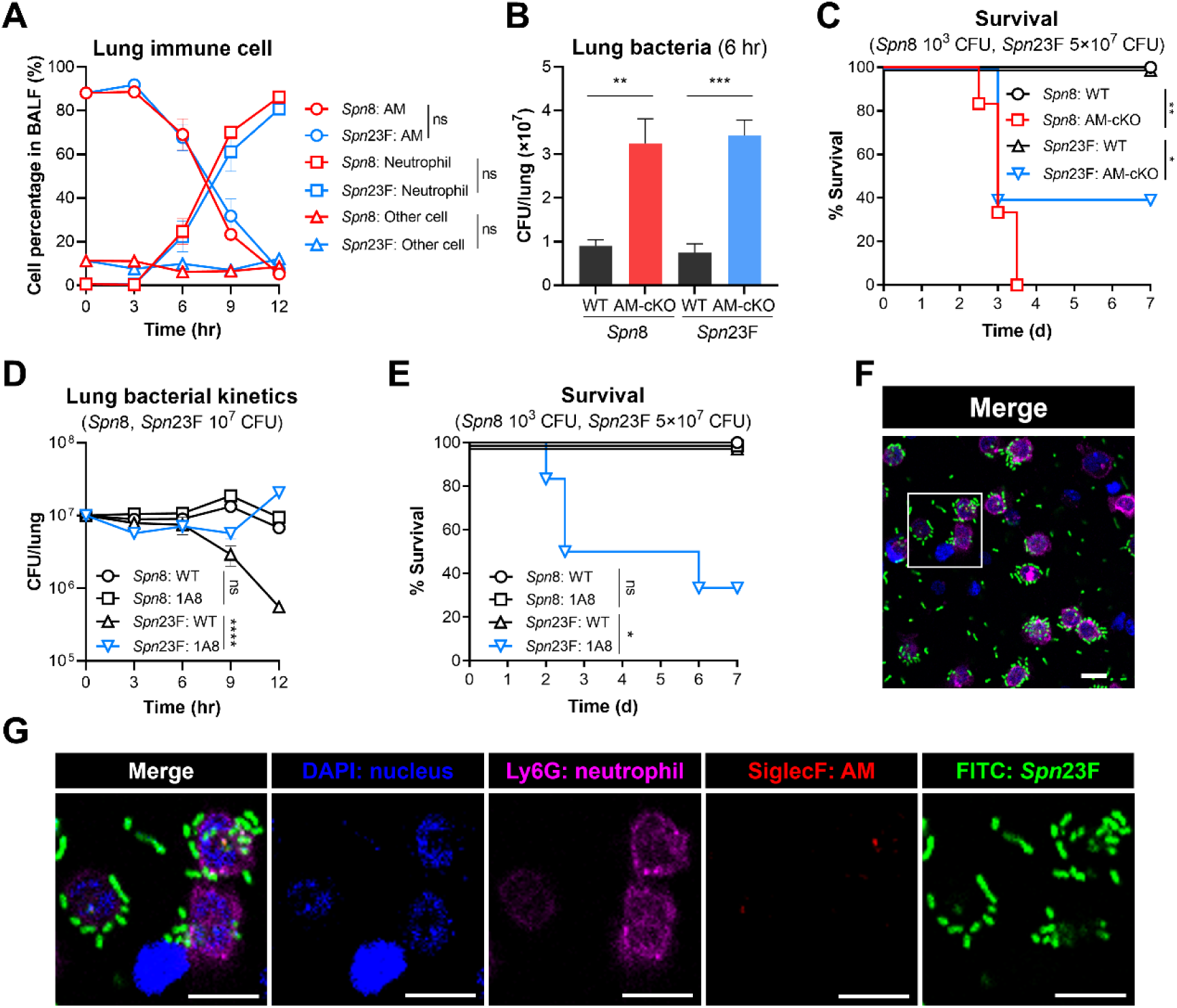
Preferential clearance of LV *S. pneumoniae* by neutrophils in the lung. **(A)** Cell percentage of AMs, neutrophils and other cells in the BALF of C57BL/6 mice i.n. infected with 10^7^ CFU of *Spn*8 and *Spn*23F. n = 3. **(B)** Bacterial loads at 6 hr in the lungs of WT or AM conditionally knockout (AM-cKO) mice i.n. infected with 10^7^ CFU of *Spn*8 and *Spn*23F. AMs were conditionally knocked out in *Cd11c*-Cre*Hdac1*^fl/fl^*Hdac2*^fl/fl^ mice. n = 3-6. **(C)** Survival time of WT or AM-cKO mice i.n. infected with 10^3^ CFU of *Spn*8 and 5 × 10^7^ CFU of *Spn*23F. n = 5-6. **(D)** Bacterial kinetics within 12 hr in the lungs of WT or neutrophil-depleted mice i.n. infected with 10^7^ CFU of *Spn*8 and *Spn*23F. Neutrophils were depleted by i.p. injection of 500 μg anti-Ly6G antibody (clone 1A8) one day before infection and daily injection of 100 μg 1A8 antibody until the end of the experiments. n = 3-6. **(E)** Survival time of WT or neutrophil-depleted mice i.n. infected with 10^3^ CFU of *Spn*8 and 5 × 10^7^ CFU of *Spn*23F. n = 5-6. **(F and G)** Fluorescent imaging (F) and enlarged area in the box (G) of *Spn*23F binding to primary neutrophils with the opsonization of normal serum. Cells were stained with DAPI to label nuclei (blue), AF647 anti-Ly6G to label neutrophils (purple), and PE anti-SiglecF to label AMs (red), simultaneously. Bacteria labeled with FITC fluorescence (green) were added at an MOI of 10. Scale bar, 10 μm. Ordinary two-way ANOVA with Sidak’s multiple comparisons test (A, D), unpaired *t* test (B), log-rank test (C, E), ns, not significant, *, P < 0.05, **, P < 0.01, ***, P < 0.001, ****, P < 0.0001.

**Figure S3.**
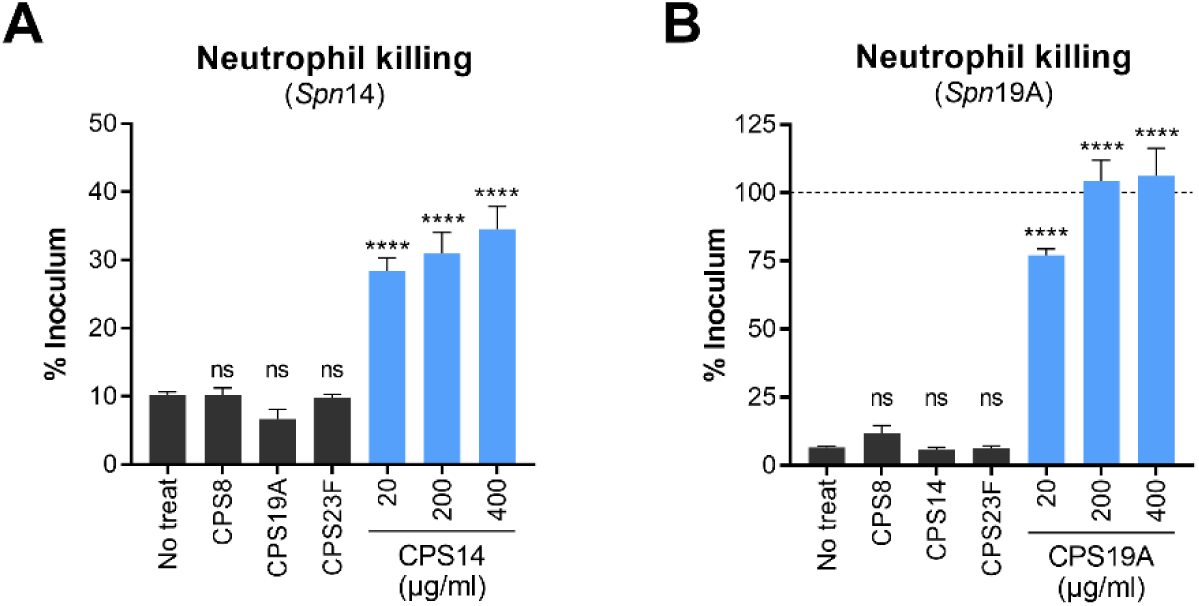
Inhibition of LV CPS to neutrophil killing of cognate LV *S. pneumoniae* serotypes. **(A)** Dose-dependent inhibition of *Spn*14 killing by primary neutrophils in the presence of free CPS14. The heterologous CPS8, CPS19A, and CPS23F were used as negative controls and added at a concentration of 400 μg/ml. n = 4. **(B)** Dose-dependent inhibition of *Spn*19A killing by primary neutrophils in the presence of free CPS19A. The heterologous CPS8, CPS14, and CPS23F were used as negative controls and added at a concentration of 400 μg/ml. n = 4. Ordinary one-way ANOVA with Dunnett’s multiple comparisons test (A, B), ns, not significant, ****, P < 0.0001.

**Figure S4.**
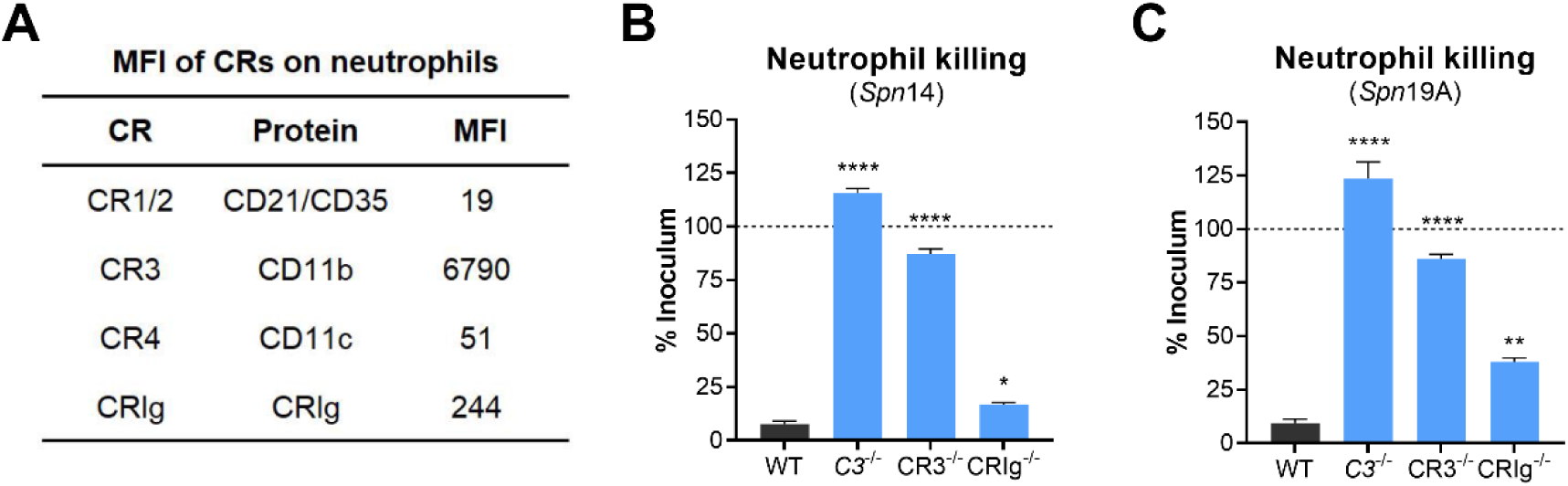
Recognition of C3-tagged LV *S. pneumoniae* by neutrophil CR3 and CRIg in lung infection. **(A)** Average MFI of CRs on the neutrophils collected at 12 hr from the BALF of C57BL/6 mice i.n. infected with 10^7^ CFU of *Spn*23F. **(B)** Killing levels of primary neutrophils to *Spn*14 isolated from WT mice with the addition of WT and *C3*^-/-^ serum, or from CR3^-/-^ and CRIg^-/-^ mice with the opsonization of WT serum. n = 4. **(C)** Killing levels of primary neutrophils to *Spn*19A under the conditions as in (B). n = 4. Ordinary one-way ANOVA with Dunnett’s multiple comparisons test (B, C), *, P < 0.05, **, P < 0.01, ****, P < 0.0001.

**Figure S5.**
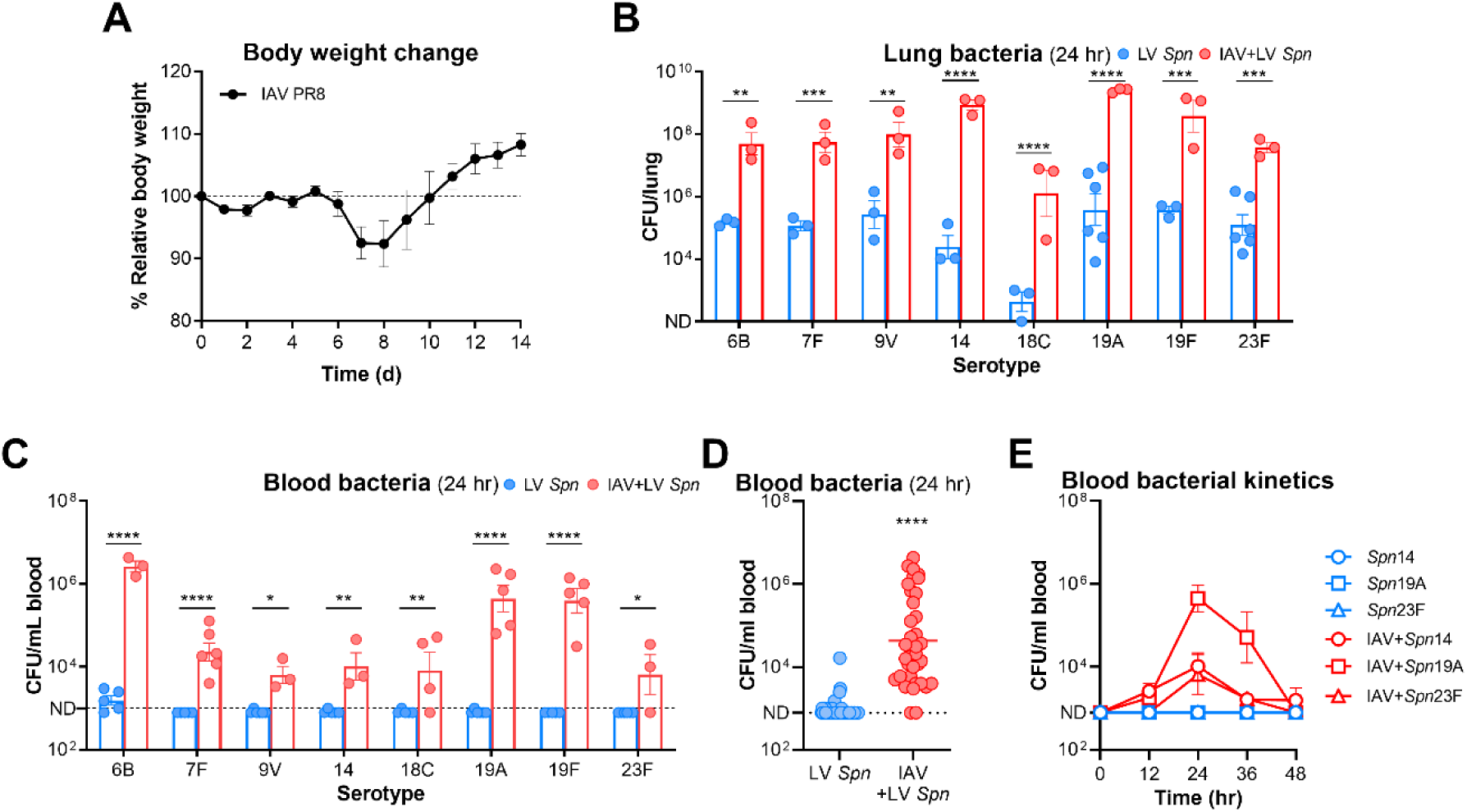
Enhanced pathogenicity of LV *S. pneumoniae* serotypes during influenza co-infection. **(A)** Body weight change of CD1 mice i.n. infected with 30 PFU of IAV PR8 strain. n = 5. **(B)** Bacterial loads at 24 hr in the lungs of mice i.n. infected with 10^7^ CFU of LV *S. pneumoniae* strains at 7 days post challenge with 30 PFU of IAV. n = 3-6. **(C and D)** Bacterial loads at 24 hr in the blood of mice infected as in (B). Data were separated for each strain (C) or combined (D). Each dot represents one mouse. n = 3-6 for each strain. **(E)** Bacterial kinetics within 48 hr in the blood of mice i.n. infected with 10^7^ CFU of representative LV *Spn*14, *Spn*19A, and *Spn*23F post IAV challenge. n = 3-6. Ordinary two-way ANOVA with Sidak’s multiple comparisons test (B, C), unpaired *t* test (D), *, P < 0.05, **, P < 0.01, ***, P < 0.001, ****, P < 0.0001.

**Figure S6.**
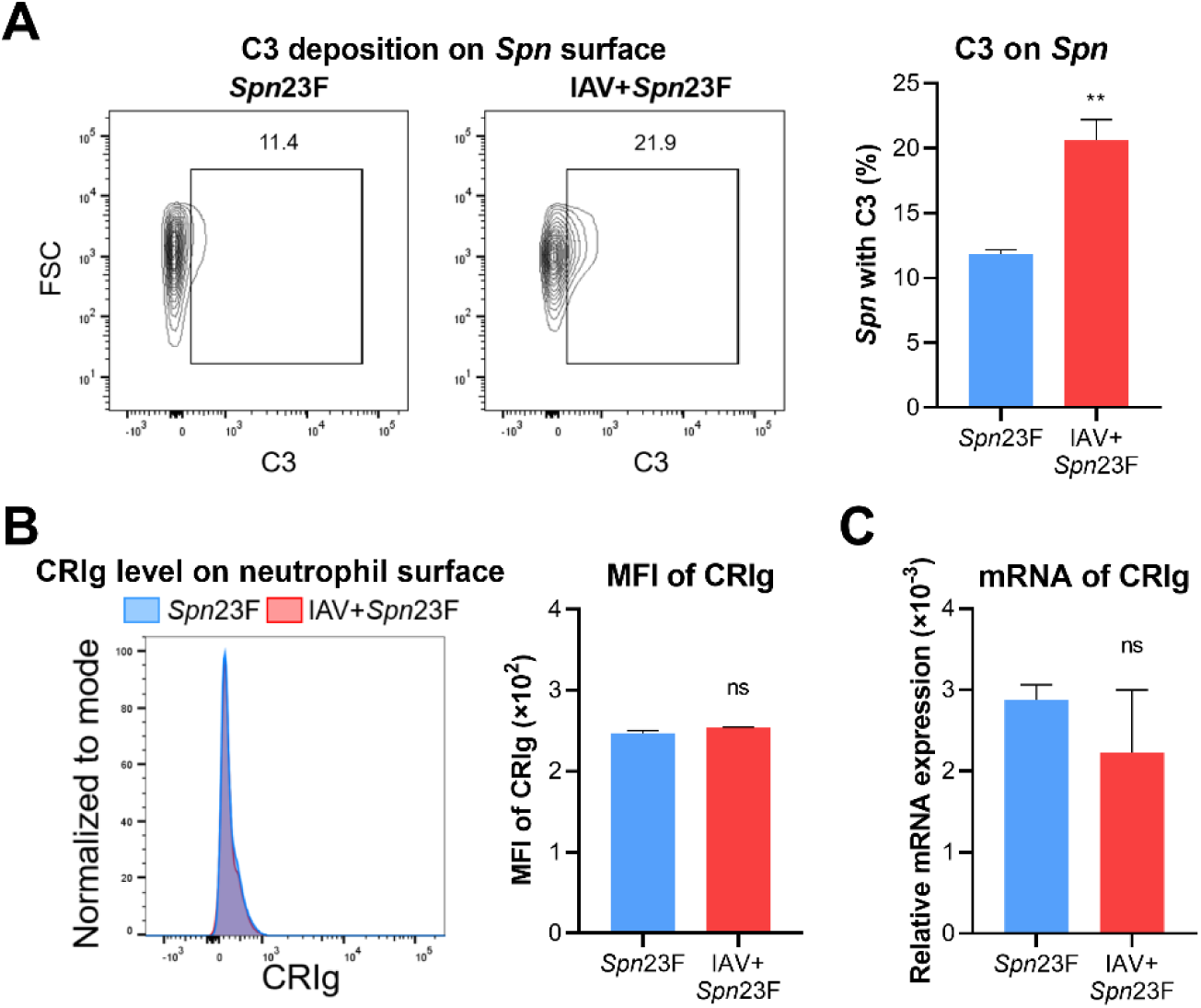
Impact of influenza infection on the C3-dependent immunity of neutrophils against LV *S. pneumoniae* serotypes. **(A)** Flow cytometry analysis of C3 deposition on *S. pneumoniae* at 9 hr collected from the BALF of C57BL/6 mice i.n. infected with 10^7^ CFU of *Spn*23F post IAV challenge. n = 3-4. **(B)** Protein level of CRIg on neutrophil surface at 12 hr isolated from the BALF of mice infected as in (A). The abundance of CRIg was determined using flow cytometry and presented as MFI. n = 3. **(C)** Transcriptional level of CRIg on neutrophils collected as in (B), and presented as the relative expression to the GADPH gene. n = 4. Unpaired *t* test (A, B, C), ns, not significant, **, P < 0.01.

**Figure S7.**
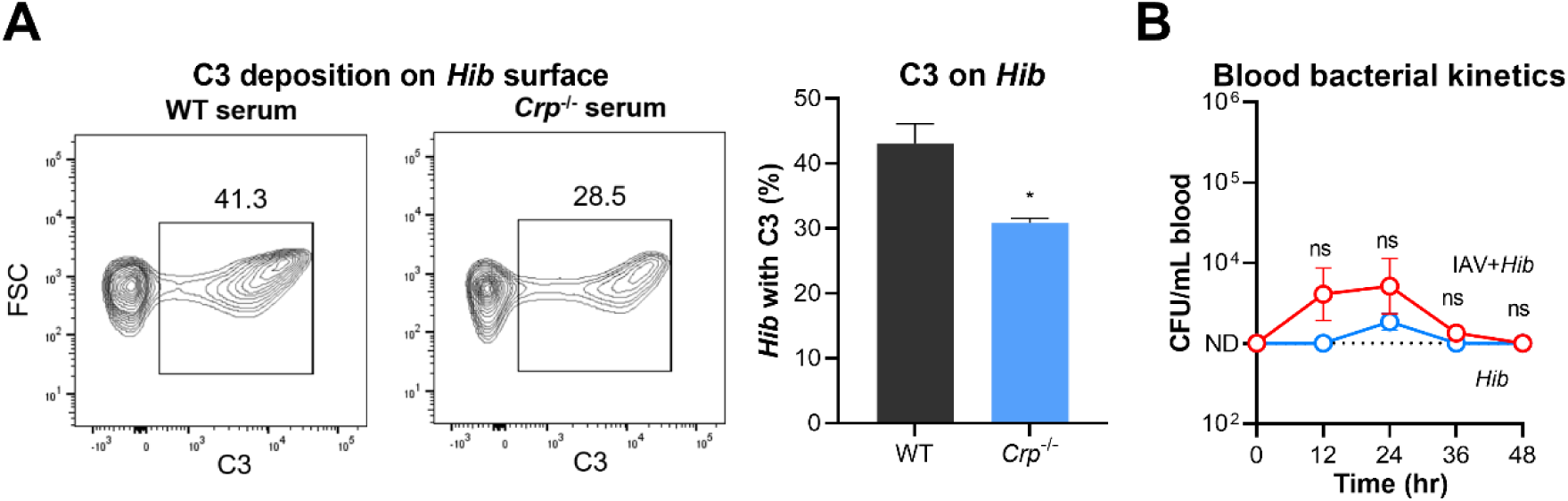
Influenza virus and *H. influenzae* synergism by suppressed CR3-mediated neutrophil immunity. **(A)** Flow cytometry analysis of C3 deposition on *Hib* surface post incubation with WT or *Crp*^-/-^ serum. n = 3. **(B)** Bacterial kinetics within 48 hr in the blood of CD1 mice i.n. infected with 5 × 10^8^ CFU of *Hib* at 7 days post challenge with 30 PFU of IAV. n = 6. Unpaired *t* test (A), ordinary two-way ANOVA with Sidak’s multiple comparisons test (B), ns, not significant, *, P < 0.05.

## SUPPLEMENTAL TABLES

**Table S1.** Information of bacterial and viral strains in this study.

| Species | Strain | Genotype | Capsule type |
| --- | --- | --- | --- |
| *S. pneumoniae* | D39 | Wild-type | 2 |
| *S. pneumoniae* | A66 | Wild-type | 3 |
| *S. pneumoniae* | TIGR4 | Wild-type | 4 |
| *S. pneumoniae* | TH870 | Wild-type | 6A |
| *S. pneumoniae* | TH2919 | Wild-type | 6B |
| *S. pneumoniae* | TH863 | Wild-type | 7F |
| *S. pneumoniae* | TH2884 | Wild-type | 8 |
| *S. pneumoniae* | TH883 | Wild-type | 9V |
| *S. pneumoniae* | TH2912 | Wild-type | 14 |
| *S. pneumoniae* | TH882 | Wild-type | 18C |
| *S. pneumoniae* | TH2918 | Wild-type | 19A |
| *S. pneumoniae* | TH2740 | Wild-type | 19F |
| *S. pneumoniae* | TH15936 | Wild-type | 23F |
| *S. pneumoniae* | TH15938 | TH870Δcps::cps2 | 2 |
| *S. pneumoniae* | TH15939 | TH870Δcps::cps3 | 3 |
| *S. pneumoniae* | TH15940 | TH870Δcps::cps4 | 4 |
| *S. pneumoniae* | TH15942 | TH870Δcps::cps6B | 6B |
| *S. pneumoniae* | TH13159 | TH870Δcps::cps7F | 7F |
| *S. pneumoniae* | TH15943 | TH870Δcps::cps8 | 8 |
| *S. pneumoniae* | TH13133 | TH870Δcps::cps9V | 9V |
| *S. pneumoniae* | TH15944 | TH870Δcps::cps14 | 14 |
| *S. pneumoniae* | TH13131 | TH870Δcps::cps18C | 18C |
| *S. pneumoniae* | TH14190 | TH870Δcps::cps19A | 19A |
| *S. pneumoniae* | TH14188 | TH870Δcps::cps19F | 19F |
| *S. pneumoniae* | TH15945 | TH870Δcps::cps23F | 23F |
| *H. influenzae* | M5216 | Wild-type | b |
| Influenza A virus | PR8 | Mouse-adapted | - |

**Table S2.**
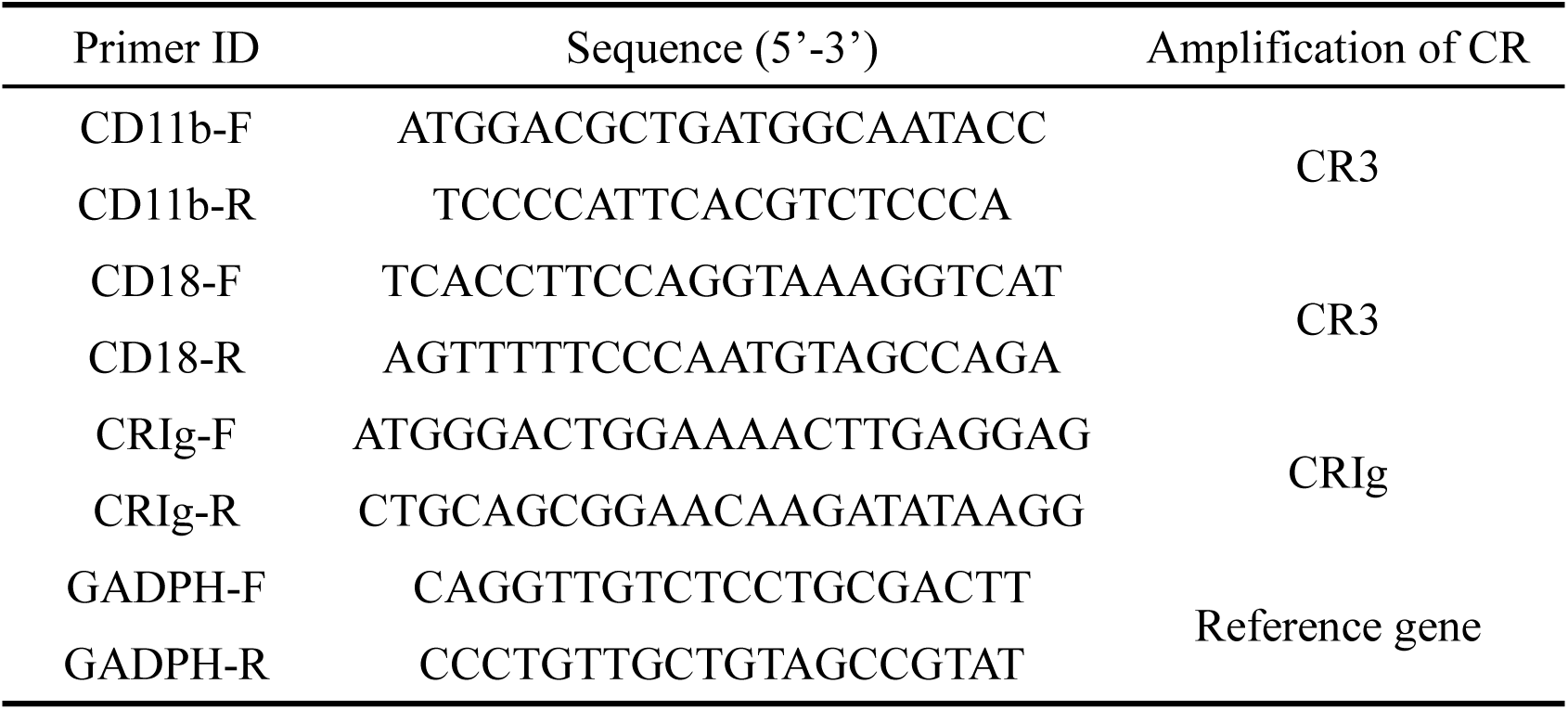
Primers used for measuring mRNA levels of CRs by qRT-PCR.

